# Crosstalk interactions between transcription factors ERRα and PPARα assist PPARα-mediated gene expression

**DOI:** 10.1101/2024.01.15.575655

**Authors:** Sofie J. Desmet, Jonathan Thommis, Tineke Vanderhaeghen, Edmee M. F. Vandenboorn, Yunkun Li, Steven Timmermans, Daria Fijalkowska, Dariusz Ratman, Evelien Van Hamme, Lode De Cauwer, Bart Staels, Luc Brunsveld, Frank Peelman, Claude Libert, Jan Tavernier, Karolien De Bosscher

## Abstract

The peroxisome proliferator-activated receptor α (PPARα) is a crucial transcription factor governing genes associated with fatty acid β-oxidation. How various interacting proteins modulate PPARα’s transcriptional function remains incompletely understood. Employing an unbiased mammalian protein-protein interaction trap with liganded PPARα as bait, we identified an interaction with the orphan nuclear receptor estrogen-related receptor α (ERRα). Random mutagenesis scanning of PPARα’s ligand-binding domain and coregulator profiling experiments implicated bridging coregulators, while *in vitro* studies suggested a trimeric interaction involving RXRα. The PPARα·ERRα interaction, dependent on three C-terminal residues within ERRα’s helix 12, was reinforced by PGC1α and serum deprivation. Pharmacological inhibition of ERRα reduced its interaction with ligand-activated PPARα, revealing a transcriptome indicative of ERRα functioning as a transcriptional repressor on prototypical PPARα target genes. Intriguingly, ERRα exhibited opposite behavior on other PPARα targets, including the isolated PDK4 enhancer. Chromatin immunoprecipitation analyses demonstrated PPARα ligand-dependent recruitment of ERRα onto specific chromatin regions where PPARα binds in mouse livers. These findings highlight intricate transcriptional crosstalk mechanisms between PPARα and ERRα, suggesting a multi-layered regulatory network fine-tuning PPARα’s activity as a nutrient-sensing transcription factor.

## Introduction

The peroxisome proliferator-activated receptor α (PPARα) and estrogen-related receptor α (ERRα) both belong to the nuclear receptor (NR) superfamily. PPARα (NR1C1) is a nutrient-sensing transcription factor involved in hepatic fatty acid (FA) transport and fatty acid β-oxidation (FAO). As such, PPARα protein levels are high in the liver, heart, skeletal muscle, or kidney. PPARs typically engage RXR partner proteins to bind cognate PPAR response elements (PPRE) of their target gene promoters ^1^. ERRs are NRs that exhibit constitutive transcriptional activity. Being so-called orphan receptors with no natural ligands identified, regulation of their expression under particular physiological or pathological conditions is an important level of control ^2^. The broadly expressed ERRα (encoded by the *ESRRA* gene) tightly controls oxidative metabolism processes in various tissues to ensure a sustained adaptive energy metabolism ^3, 4^. For example, ERRα regulates hepatic gene expression by exerting opposing effects on genes important for mitochondrial oxidative capacity and gluconeogenesis ^3, 5^. XCT790 or C29 are synthetic inverse agonists that can suppress ERRα’s constitutive activity and some of these pharmacological agents were shown to be of benefit in diabetic mouse models ^6^.

The transcriptional activity of ERRα is typically regulated by coregulators such as PPARγ coactivator-1 alpha (PGC-1α) ^5^, recently shown to functionally connect ERRα with transcriptional complex components to overcome promoter-proximal pausing of RNA polymerase (pol) II ^7^, but also by posttranslational mechanisms including acetylation and phosphorylation ^2, 8^. Stimuli that increase FAO, such as fasting, increase the expression and activity of PPARα, ERRα, and their common coactivator PGC-1α in the liver ^1, 3, 9^. A close functional connection between both receptors is illustrated by the observation that many ERRα-regulated genes include targets of PPARα. This is further supported by the finding that ERRα binding onto the promoter of PPARα activates *Ppara* gene expression in mouse myocytes ^10^. Another key common target gene of both PPARα and ERRα in liver is *pyruvate dehydrogenase kinase 4* (*PDK4*) ^11, 12^. The PDK4 enzyme blocks mitochondrial pyruvate oxidation in favor of FAO and hence serves as a crucial checkpoint between glucose and lipid metabolic pathways^12–14^.

Here, we studied the molecular determinants explaining transcriptional crosstalk mechanisms between PPARα and ERRα-regulated pathways in various cellular models and starved murine livers. Fasting was chosen as a stimulus known to induce *ESRRA* gene induction and as such ERRα activity in hepatocytes. Pharmacological hampering of ERRα activity positively or negatively affected PPARα transcriptional activity depending on the nature and regulatory elements in the vicinity of the target gene. Upon retrieval of ERRα among top candidates in a cell-based screen for PPARα interacting proteins, we studied whether both proteins may reside within the same complex. Shifts in cofactor recruitment profiles may be contributory mechanisms to explain gene expression changes of particular gene targets under transcriptional co-control of PPARα and ERRα, e.g. *PDK4*. Comparative mouse fasted-fed liver studies support the existence of a coordinate nuclear receptor crosstalk mechanism which is surprisingly more outspoken in the fed state. Following PPARα agonist treatment, ERRα is recruited at the chromatin of known PPARα-controlled promoters and enhancer DNA in fed livers. The marked PPARα agonist-induced ERRα recruitment in fed livers is lost again following ERRα inhibition.

## Results

### Ligand-activated PPARα interacts with ERRα more strongly within cells than in vitro

The mammalian two-hybrid technology MAPPIT, for Mammalian Protein-Protein Interaction Trap, entails a cell-based screening system to identify interaction partners of a given bait protein. Only an interaction between ‘bait protein’ and ‘prey protein’ brings functional cytokine signalling components together and restores a leptin-inducible cytokine receptor signalling pathway followed by a STAT3-dependent luciferase read-out. The system is highly sensitive and designed to also detect transient interactions ^15, 16^. We applied MAPPIT with the PPARα-specific agonist GW7647 (GW)-liganded PPARα as bait, against a human ORFeome collection of 8.500 preys. ERRα was identified as a top candidate interactor of ligand-activated PPARα, among well-known direct PPAR family interactors RXR ^17, 18^ and NR0B1 ^19^ (Fig. 1A; Supplementary Fig. S1A). Independent binary MAPPIT re-tests (Fig. 1B) and co-immunoprecipitation analysis (Fig. 1C) confirmed a GW-induced interaction between ERRα and PPARα. The MAPPIT technology relies on overexpression and constrains PPARα to the membrane. Hence, to localize both NRs within the cell, we used primary murine hepatocytes given they contain high levels of endogenous PPARα and ERRα. Immunofluorescence analysis (Fig. 1D) confirms earlier reports that both endogenous NRs mainly reside in the hepatocyte nucleus, even in the absence of any (synthetic) ligands ^20, 21^. In the presence of GW, added for 1h, colocalization as indicated by the Pearson’s correlation coefficient, is significantly enhanced (Fig. 1D). A similar nuclear overlay profile is also observed for the human hepatocyte cell line HepG2 (Supplementary Fig. S1B). A proximity ligation assay in GW-induced HepG2 revealed a dotted signal which suggests the possibility of an endogenous interaction between PPARα and ERRα (Figure 1E). Complementary *in vitro* GST-pulldowns (Fig. 1F,G; Supplementary Fig. S1C) support that full-length PPARα and ERRα may interact, either directly or indirectly, while His-pulldown assays suggest that PPARα’s ligand binding domain (LBD) and full-length ERRα can interact directly, albeit weakly (Fig. 1H; Supplementary Fig. S1D). In line with weak binding, or, reflecting the need for a stabilizing factor within cells, ERRα was unable to displace Med1 or p300 from PPARα in an *in vitro* competition assay (data not shown). Results using *in vitro* transcribed and translated ERRα from wheat germ extract to exclude the presence of mammalian coregulators (Fig. 1F), reliably mirrored PPARα interaction with ERRα protein from reticulocyte lysate (Supplementary Fig. S1C). A GST fusion of the coactivator protein PGC1α, a known direct ERRα-interactor ^22^, served as positive control. Because PPARα heterodimerizes with RXR family members, we investigated the impact of additional ERRα on the PPARα-RXRα complex, via GST-pulldown assays with GST-RXRα (or its DNA-binding domain as a negative control). Fig. 1G shows that full-length RXRα (lane 3), interacts with PPARα, as expected. With increasing amounts of ERRα protein more PPARα is pulled down, suggesting that complex formation with all three proteins is possible (lane 4). Titrating in more ERRα maintains the complex, yet, less PPARα is pulled down with RXRα. Contrasting to the cell-based data, the *in vitro* interaction studies did not support GW ligand-dependency (Fig. 1F, Supplementary Fig. S1C). Nevertheless, two different ERRα inverse agonists, C29 and XCT790, did lower the observed interaction between PPARα-LBD and ERRα (Fig. 1H). A control fluorescence polarization assay in presence of XCT790 (Supplementary Fig. S1E) confirms that the recombinant ERRα protein used in the His-pulldown assays is functional. Collectively, even though a direct interaction between PPARα and ERRα is GW ligand-independent or weak *in vitro*, a much stronger GW ligand-dependent and potentially indirect interaction, can be observed within cells.

**Fig. 1.**
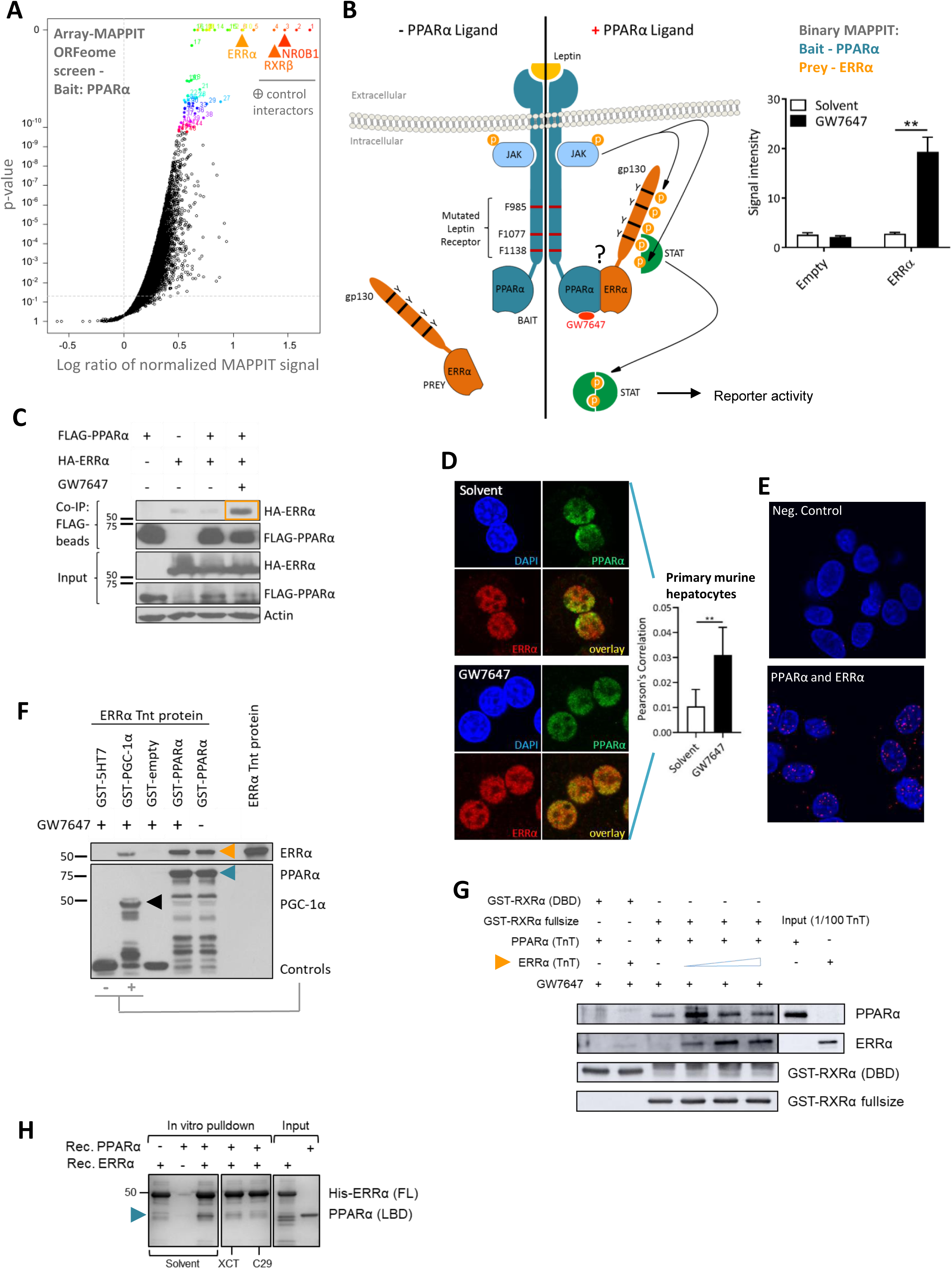
ERRα can interact with ligand-activated PPARα in vitro and in cellulo. (A) Array mammalian protein-protein interaction trap (MAPPIT) screen result of the activated (with agonist GW7647) PPARα bait against a prey library shown as a volcano plot displaying the log ratio of normalized MAPPIT luciferase activity versus *p*-value (horizontal line: p-value= 0.05). (B) Binary MAPPIT principle and re-test result with PPARα as bait and ERRα as prey (empty prey: neg. ctrl). Cells were stimulated with leptin (100ng/ml) +/-GW7647 (0.5µM) for 24h or were left untreated. Luciferase measurements were normalized by untreated values (n=5, mean + SEM). (C) Representative co-immunoprecipitation between Flag-PPARα and HA-ERRα, overexpressed in HEK293T cells (GW7647 stimulation: 0.5µM, 3h) (n=3). (D) Confocal immunofluorescence results of GW7647 (0.5µM, 1h)-stimulated primary murine hepatocytes for PPARα (green) and ERRα (red); nuclei were visualized with DAPI staining (blue) and thresholded Pearson’s correlation coefficients were calculated after scanning (mean + SD, unpaired t-test). (E) Endogenous interaction analysis of PPARα and ERRα on GW7647 (0.5µM, 1h)-stimulated HepG2 cells via proximity ligation assay (PLA). Cells were stained with mouse anti-PPARα and rabbit anti-ERRα antibody (nuclei stained with DAPI (blue)), followed by an anti-mouse MINUS and anti-rabbit PLUS probe, respectively. Positive signal (red dots) is generated when proteins of interest are within 40nm. Depicted are merged images including the negative control without primary antibodies (top). (F) GST-pulldown between GST-PPARα and rabbit reticulocyte *in vitro* transcribed and translated ERRα (ERRα TnT protein) (GST-5HT7, GST-empty: neg. ctrls; GST-PGC-1α: pos. ctrl; GW7647 at 5µM). (G) GST-pulldown analysis to probe the interaction of GST-RXRα fusion proteins with *in vitro* transcribed and translated PPARα and ERRα (n=2). A fixed amount of GW7647-activated PPARα in the absence or presence of increasing amounts of ERRα was incubated with glutathione-agarose 4B beads loaded with GST-RXRα (DBD) (negative control) or GST-RXRα full-size protein. (H) In vitro His-tag pulldown of PPARα-LBD and full-length His-ERRα in the presence of the ERRα inhibitors XCT790 and C29 (n=3)

### Pharmacological inhibition of ERRα and mutagenesis at its C-terminus abrogates the functional interaction with PPARα

Because PPARα is a sensor of nutrient-deprivation, we wondered whether the absence of serum in the cell culture could affect the interaction. Serum starvation enhanced the MAPPIT signal with a factor of almost 3 (Fig. 2A). To exclude that the effect of GW might be compound-specific, we next included Pemafibrate (Pema), another agonist of PPARα ^23^. Pema supported the interaction between PPARα and ERRα equally well as GW (Fig. 2B) and both MAPPIT interaction profiles were efficiently blocked with C29, a newer-generation pharmacological inhibitor of ERRα ^13, 24, 25^. To investigate the impact of ERRα on the inherent transcriptional capacity of PPARα, independent of DNA binding events, we used a mammalian one-hybrid transcriptional system consisting of a Gal4 DBD coupled to PPARα and a Gal4-dependent reporter gene (Fig. 2C). Exogenous ERRα enhanced the transcriptional activity of both pema- and GW-activated Gal4-PPARα (Fig. 2C). Blocking ERRα with C29 efficiently diminished the ligand-induced transcriptional activity of Gal4-PPARα. ERRα expression controls demonstrate that the reduced transcriptional activity in presence of C29 was not due to less ERRα protein (Supplementary Fig. S2A). Using a Gal4-PPARα LBD fusion protein indicated that the interaction involves the LBD of PPARα (Supplementary Fig. S2B). Protein localization experiments verified that a combined C29/GW treatment kept both proteins within the nucleus (Supplementary Fig. S2C). Collectively, so far the data suggest that a nuclear interaction between PPARα and ERRα can occur via the LBD of PPARα and that the interaction can be enhanced in serum-deprived cells. To study which amino acids of the PPARα-LBD protein surface are important for the interaction with ERRα, we coupled an extensive random mutagenesis screen to MAPPIT. Error prone PCR to randomly mutate PPARα-LBD yielded an amino acid coverage of 56%. We cloned single (and some selected double)-mutants as full-length PPARα in bait plasmids and screened for interactions with ERRα in the presence of GW. When mapping the interaction results onto the PPARα-LBD crystal structure, a discrete ERRα-interaction sensitive binding region emerged (Fig. 2D, red residues). This surface co-incidentally corresponds to the PPARα coactivator binding site. Interaction-dead mutants (red), independently validated by conventional MAPPIT (Supplementary Fig. S2D), expressed equally well as wildtype (WT) PPARα (Supplementary Fig. S2E) while three out of seven mutations had no impact on protein stability, as predicted by FoldX calculations (Supplementary Fig. S2F). In line with the PPARα agonist-dependency of the interaction *in cellulo* (Fig. 1A-C, Fig. 2A-C), the confined binding surface shifts to a more scattered profile when overlaid on the PPARα-LBD structure in antagonist instead of agonist mode (Fig. 2E). Upon considering the existence of a direct interaction and taking the identified interaction surface into account, molecular modelling predicts that helix 12 (H12) of ERRα may directly bind onto the PPARα coactivator site (Supplementary Fig. S2G). To test this putative interaction surface, we mutated three hydrophobic amino acids in ERRα H12 to alanine (A), more specifically the methionine (M) at position 417 and 421, and the leucine (L) at position 418 (Fig. 2F). MAPPIT revealed that the resulting ERRαMLM mutant no longer interacts with the activated PPARα bait, in sharp contrast to WT ERRα (Fig. 2F). In line, the ERRαMLM mutant also fails to support GW-induced Gal4-PPARα activity (Fig. 2G). We verified that both full-length and the ERRαMLM mutants were highly expressed (Fig. 2F and 2G). Taken together, the tail of ERRα’s helix 12 is important for the interaction with PPARα.

**Fig. 2.**
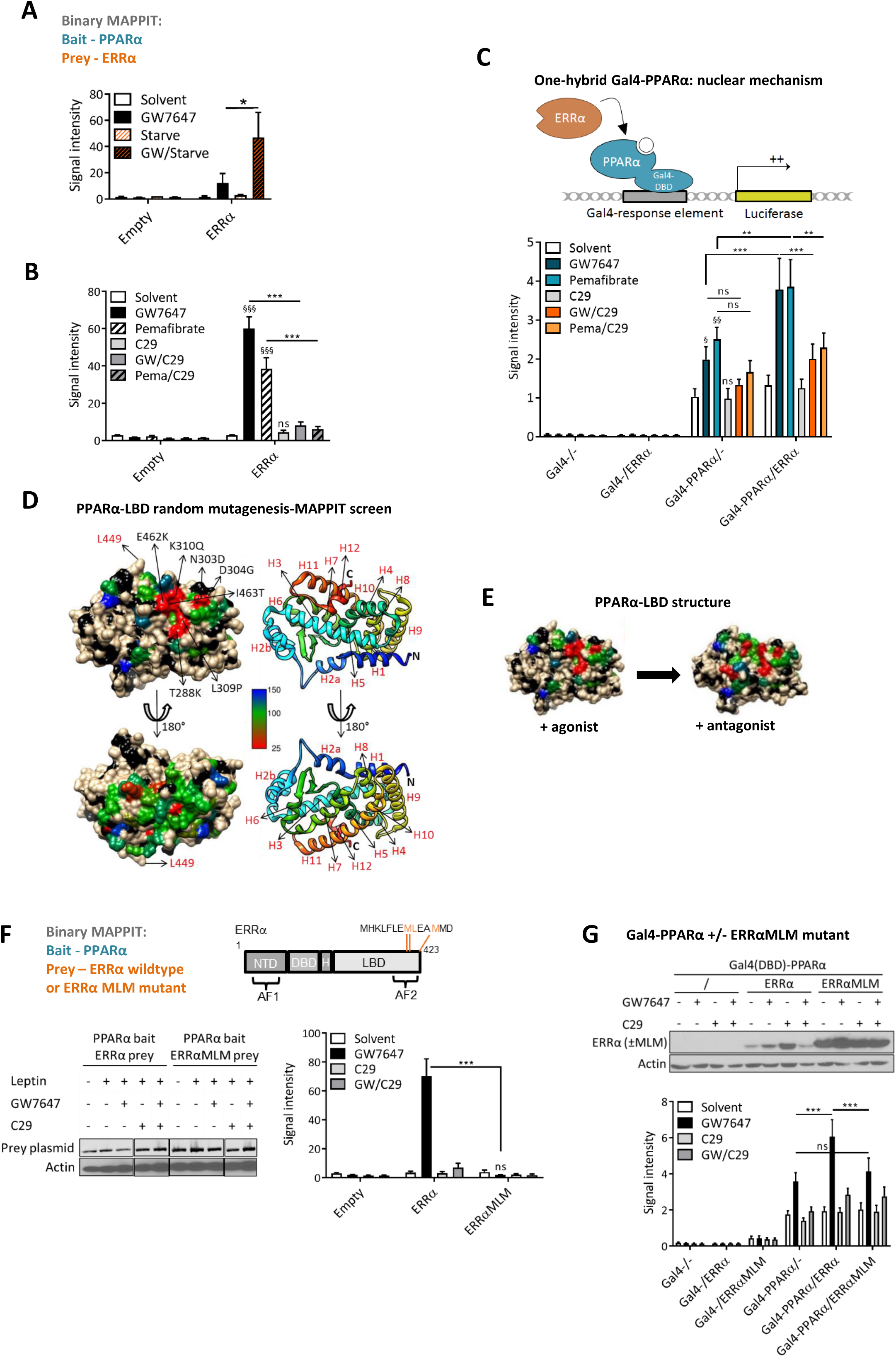
Pharmacological inhibition of ERRα and mutagenesis at its C-terminus abrogates the functional interaction with PPARα. (A) Binary MAPPIT with PPARα as bait and ERRα as prey (empty prey: neg. ctrl). HEK293T cells were stimulated with leptin (100ng/ml) +/- GW7647 (0.5µM) for 24h or were left untreated. Cell stimulations were either in the absence (“starve”) or presence of serum in the culture media. Luciferase measurements were normalized by untreated values (n=3, mean + SEM). (B) Binary MAPPIT with PPARα as bait and ERRα as prey (empty prey: neg. ctrl). Cells were serum-starved, stimulated with leptin, induced +/- GW7647 (0.5µM) or pemafibrate (5µM) and/or the ERRα inverse agonist C29 (5µM) for 24h, or were left untreated. Luciferase measurements were normalized by untreated values (Mean + SEM, n=3). In the ERRα prey condition, the significance of differences in binding activity was evaluated with one-way ANOVA, followed by multiple comparison using the Fisher’s LSD test (*: p<0.05; **: p<0.01; ***: p<0.001, significance of single compound vs Solvent is marked with § signs). (C) Principle of mammalian one-hybrid experiment. PPARα full-length is coupled to the DNA binding domain of Gal4 (“Gal4-“), which can bind its response element and activate the luciferase reporter. HepG2 cells, transfected with Gal4-responsive luciferase reporter, Gal4- (control) or Gal4-PPARα, and with or without ERRα, were stimulated with GW7647 (GW, 0.5µM) and/or C29 (5µM) for 24h or were left untreated (Mean + SEM, n=3). The significance of differences in reporter activity was evaluated with unbalanced two-way ANOVA, followed by multiple comparisons using the Fisher’s LSD test (*: p<0.05; **: p<0.01; ***: p<0.001, significance of single compound vs Solvent is marked with § signs). (D) Random mutagenesis-interaction result on the PPARα-LBD crystal structure (PDB: 2P54) is shown in two different orientations (residue L449 is marked as a reference point (red font)). On the surface presentation, the color of the residues corresponds to the relative MAPPIT signals (% of wild-type), and thus binding effect, and ranges from red (<25%) to blue (>150%) (backbone atoms: black, side-chains of non-mutated residues: white). (E) Comparison of PPARα-LBD mutagenesis results when modelled in complex with an agonist (PDB: 2P54) versus an antagonist (PDB: KKQ) (F) MAPPIT with PPARα as bait and ERRα or ERRαMLM mutant as prey (empty prey: neg. ctrl). Serum-starved cells were stimulated with leptin +/- GW7647 (0.5µM) and/or C29 (5µM) for 24h or were left untreated. Luciferase measurements were normalized by untreated values (Mean + SEM, n=3). In the ERRα and ERRαMLM prey conditions, the significance of differences in binding activity was evaluated with two-way ANOVA, followed by multiple comparison using the Fisher’s LSD test (*: p<0.05; **: p<0.01; ***: p<0.001). Corresponding protein expression controls of mutant and wildtype ERRα protein are depicted. (G) HepG2 cells, transfected with Gal4-responsive luciferase reporter, Gal4(DBD)-control or Gal4(DBD)-PPARα, and with or without ERRα or the triple mutant ERRα (ERRαMLM), were stimulated with GW7647 (GW, 0.5µM) and/or C29 (5µM) for 24h or were left untreated (Mean + SEM, n=3). In the Gal4-PPARα conditions, the significance of differences in reporter activity was evaluated with unbalanced two-way ANOVA, followed by multiple comparison using the Fisher’s LSD test (*: p<0.05; **: p<0.01; ***: p<0.001). Western analysis depicts the corresponding loading controls.

### The interaction between PPARα and ERRα is strengthened by PGC1α

One way to explain the discrepancy between weaker *in vitro* and much stronger *in cellulo* binding results could be the presence of a PPARα ligand-responsive, interaction-bridging, coregulator in the cells. A logical candidate to investigate is PGC1α, a coregulator known to bind both PPARα and ERRα and given a role in past literature as the protein-ligand of ERRα^10, 26^. Conveniently, different mutants of PGC1α have been described (Fig. 3A), able to interact exclusively to ERRα (L2A mutant), retaining the ability to interact with both receptors (L3A) or losing the interacting capacity to both receptors (L2A/L3A). We monitored the impact of PGC1α WT and mutants on the binary interaction between ligand-activated PPARα and ERRα via MAPPIT. Surprisingly, not only PGC1α WT, but also overexpression of the L2A mutant (which can only interact with ERRα) could enhance the MAPPIT interaction between GW-activated PPARα and ERRα, while the L2/L3A mutant still allowed for a residual basal interaction profile (Fig. 3A). The data seem to suggest that endogenous PGC1α or another coregulator may be sufficient to support a basal interaction profile and that a strengthened interaction axis between PGC1α and ERRα enhances the latter receptor’s interaction capability with PPARα.

**Fig. 3.**
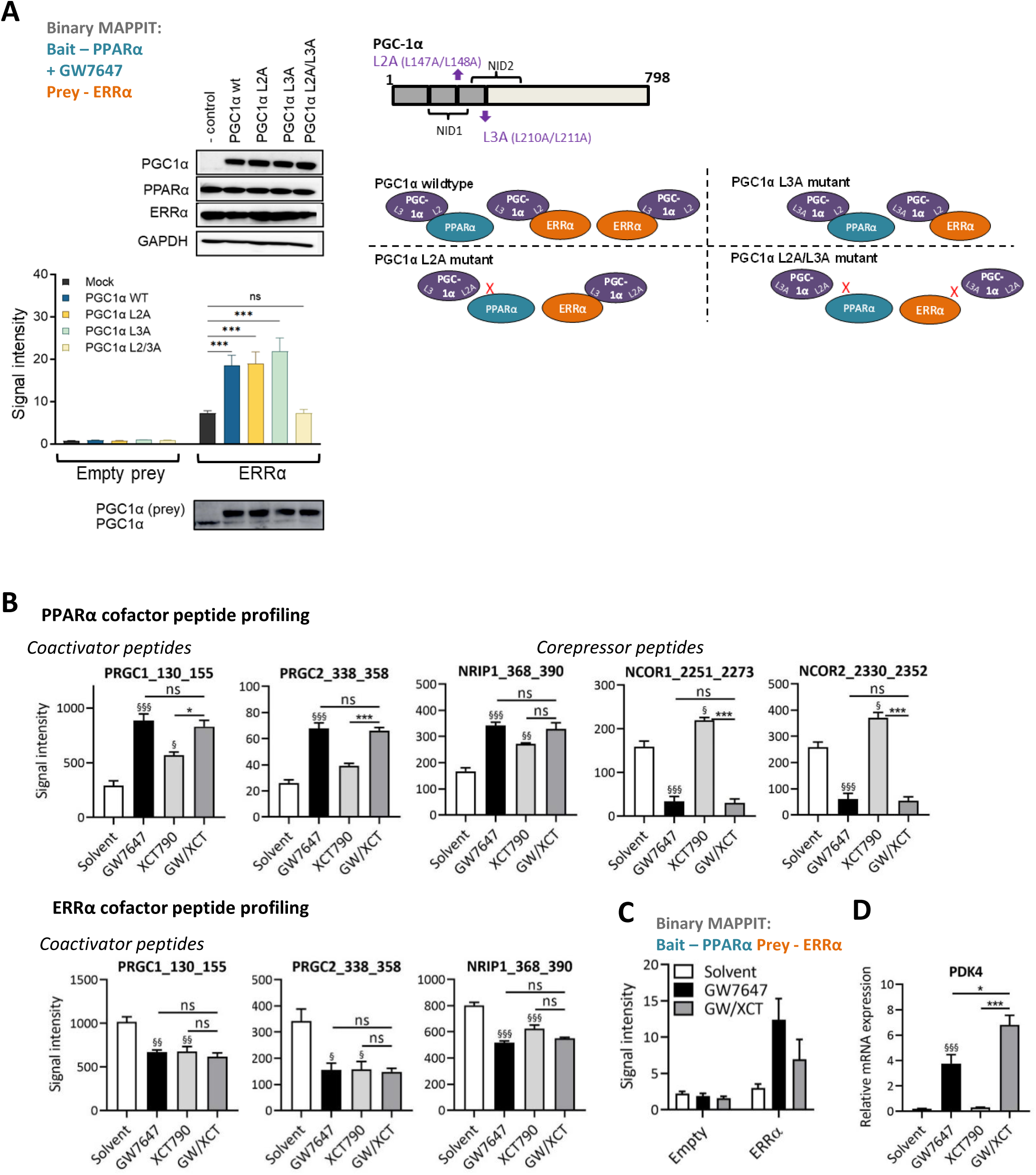
The interaction between PPARα and ERRα is strengthened by ERRα’s protein ligand PGC1α and PPARα agonism decreases ERRα’s constitutive coregulator recruitment profile. (A) MAPPIT with PPARα as bait, ERRα as prey (empty prey: neg. ctrl) and supplemented with PGC1α WT or mutants as indicated on the protein domain figure. Serum-starved HEK293T cells were stimulated with leptin + GW7647 (0.5µM) throughout. Corresponding protein expression controls are depicted (top) while a reload blot (bottom) indicates endogenous PGC1α levels. Expected compromised interactions between NRs and PGC1α mutants conform literature are indicated in the graphic model. Luciferase measurements were normalized by untreated values (Mean + SD). (B) MARCoNI assay results (full panel, see Supplementary Fig. S3a) with in the top panel HA-PPARα, in the presence of FLAG-ERRα, overexpressed in HEK293T cells, serum-starved and stimulated with GW7647 (0.5µM) and/or XCT790 (1µM) for 2h. The lower panel shows the result of the MARCoNI assay (full panel, see Supplementary Fig. S3B) with HA-ERRα, in the presence of FLAG-PPARα (lower panel), overexpressed in HEK293T cells, serum-starved and stimulated with GW7647 (0.5µM) and/or XCT790 (1µM) for 2h. Data was fitted according to a LOESS regression (Mean + SEM, n=3). Signal intensity corresponds to the binding strength of the respective receptors to the immobilized peptides derived from PGC1α/β (=PRGC1/2), NCOR1/2 (for PPARα) and NRIP1. For each peptide, the significance of differences in binding activity was evaluated with one-way ANOVA, followed by multiple comparisons to the reference level GW/XCT using the Sidak t-test. *: p<0.05; **: p<0.01; ***: p<0.001, the significance of single compound vs Solvent is marked with § signs. (C) MAPPIT with PPARα as bait and ERRα as prey (empty prey: neg. ctrl). Serum-starved cells were stimulated with leptin +/- GW7647 (0.5µM) and/or the ERRα inverse agonist XCT790 (1µM) for 24h or were left untreated. Luciferase measurements were normalized by untreated values (Mean + SD). (D) Serum-starved HepG2 cells were stimulated with GW7647 (0.5μM) and/or the ERRα inverse agonist XCT790 (1μM) for 24h. RNA expression values of the PDK4 gene were normalized to the reference genes GAPDH and TBP using qBase+. Means + SEM (n=4) are shown on the original scale. The significance of differences in ligand effects (on the log-transformed scale) was evaluated with one-way ANOVA (***), followed by multiple comparisons to the reference level GW/XCT using the Sidak t-test.

### PPARα agonism decreases ERRα’s constitutive coregulator recruitment profile

To further investigate the possibility of a PPARα ligand-dependent NR-bridging coregulator constellation or competition mechanisms, we next studied within the same cell system both receptors’ coregulator recruitment profiles via the peptide Micro-Array for Real-time Coregulator and Nuclear receptor Interactions (MARCoNI) assay ^18^ (Fig. 3B). The MARCoNI assay was thus performed using lysates from differently tagged receptor-transfected HEK293T cells. As expected for PPARα, GW supported significant recruitment of PGC1α/β (PRGC1/2) and NRIP1 coactivator peptides, and a concomitant loss of corepressor NcoR1/2 peptides (Fig. 3B, top panel). Interestingly, XCT790 alone supported both a slightly enhanced coactivator as well as NcoR1/2 corepressor recruitment to PPARα, suggesting ERRα might influence PPARα-coregulator equilibria. There were no differences however between GW/XCT and GW, which suggests that either a plateau was reached or that the GW effect may be dominant (Fig. 3B, top panel). Oppositely and most intriguingly, constitutive PRGC1/2 and NRIP1 recruitment were consistently diminished when monitoring the cofactor recruitment profile of ERRα for all ligands separate or combined (Fig. 3B, bottom panel). Corepressors were not detected with ERRα (Supplementary Fig. S3 depicts the full panels). The GW/XCT coregulator profile of ERRα follows the GW-alone or XCT-alone profiles. Just as for C29 (Fig. 2B), MAPPIT analysis verified also XCT790’s ability to diminish the interaction between ERRα and PPARα (Fig. 3C). Downstream of changes in coregulator interaction profiles, we next asked whether XCT790 may also influence GW-induced PPARα-mediated gene transcription. mRNA levels of a prototypical PPARα target gene, *PDK4*, were enhanced when combining GW with XCT790 versus GW alone, which is suggestive of a role for ERRα as an inhibitor of this target (Fig. 3D). Collectively, the data are in support of a transcriptional crosstalk mechanism between PPARα and ERRα, with GW-mediated ERRα coactivator losses as most remarkable and unexpected features.

### Liver transcriptome analysis shows enhanced PPARα-mediated fatty acid degradation and oxidative phosphorylation processes when inhibiting ERRα

Under caloric restriction, liver PGC1α is well documented to be upregulated ^27, 28^. In addition, we previously showed that PPARα target genes in mouse livers can be further increased by GW, following overnight fasting ^29^. Given the interaction between PPARα and ERRα can be strengthened by PGC1α (Fig. 3A), we opted to first investigate a potential transcriptional crosstalk in murine livers in a context of food deprivation. Fasted mice were injected (i.p.) with GW and/or C29 and sacrificed 4h later (Fig. 4A) followed by RNAseq analysis (3 mice/group) (Supplementary Table 1 for the full RNAseq dataset). Plots of normalized reads of genes coding for PPARα, ERRα and for the downstream target PDK4 indicate ligand responsiveness *in vivo* (Fig. 4A) and a similar regulation of *Pdk4* as in HepG2 (Fig. 3D). Hierarchical data clustering supports again a dominant effect of the GW ligand with only modest changes when combined with C29 (Supplementary Fig. S4A). A fairly large overlap between GW and C29 plus GW gene clusters is apparent, as shown by the venn diagram (Fig. 4B). Therefore, we limited the subsequent analysis to address whether or not a unique effect is observed only upon combined ligand treatment (Fig. 4C-D). As such, gene set enrichment analysis of GO terms and KEGG pathways in the interaction analysis revealed a significant difference, i.e. an enrichment at the front of a gene list ordered based on log2FC attributed to interaction, a.o. for both mmu00071 (FA degradation) and mmu00190 (Oxidative phosphorylation) pathways (Fig. 4C). Network analysis confirmed FA metabolism, oxidative phosphorylation and the PPAR signaling pathway (Fig. 4D). Even though effect sizes are small, causing an average expression increase of 18% in RNAseq, GW/C29 consistently upregulates mRNA for a subset of targets when compared to GW alone, as is the case for *Pdk4* (Fig. 4A, Supplementary Fig. S4B). Because the interaction analysis pointed to changes in the oxidative phosphorylation pathway, causing on average a 9% upregulation, normalized reads of some components hereof are plotted (Supplementary Fig. S4C). To validate the trends and increase power, we next analyzed RT-qPCR transcripts from all livers (= 6 mice/group) (Fig. 4E), and confirmed the modest yet consistent transcriptional upregulation on key PPARα target genes involved in fatty acid metabolism *(Pdk4, Cpt1/2, Acaa1b, Ehhadh, Lpl*) and on the CPT1α protein level (Fig. 4F). We recapitulated similar regulations and combined ligand crosstalk in serum-starved HepG2, both at the transcript level and the CPT1α protein level (Supplementary Fig. S4D, Fig. S4E). So far, our data suggest that ERRα can behave as a subtle transcriptional repressor of PPARα-driven genes involved in fatty acid metabolism.

**Fig. 4.**
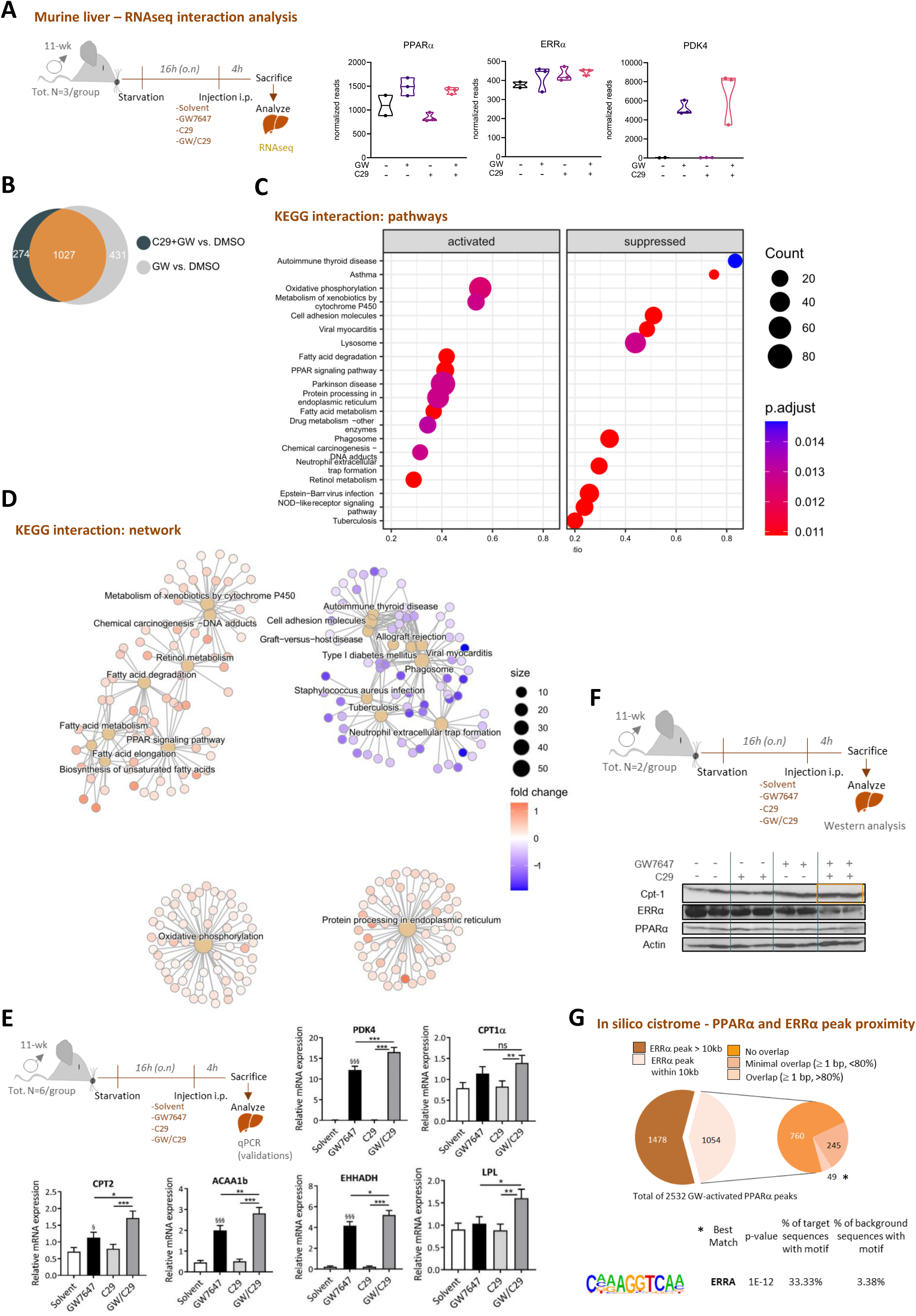
Liver transcriptome analysis shows ERRα’s ability to mitigate PPARα-mediated fatty acid degradation and oxidative phosphorylation processes. (A) C57BL/6J mice, after 16h starvation, were treated for 4h with GW7647 (4mg/kg) and/or C29 (10mg/kg) or were given solvent control, via i.p. injection. Livers were isolated for RNAseq analysis. Normalized reads of data from a pilot RNAseq experiment (3 mice/group) are shown for GW7647- regulated transcripts *Pdk4, Ucqr10, Ppara* and *Esrra*. (B) The Venn diagram shows vast overlap between the gene cluster regulated by GW7647 (vs Solvent) and GW/C29 (vs Solvent). KEGG pathways (C) and networks (D) enriched upon GW and C29 interaction are visualized using enrichplot and pathview. (E) As an independent validation, C57BL/6J mice, after 16h starvation, were treated for 4h with GW7647 (4mg/kg) and/or C29 (10mg/kg) or were given solvent control, via i.p. injection. Livers (6 mice/group) were analyzed via qRT-PCR. RNA expression values were normalized to the reference genes *Cyclo* and *b-actin* using qBase+. The significance of gene-specific ligand effects, estimated as the difference (on the transformed scale) to the gene-specific reference level GW/C29, was assessed (*: p<0.05; **: p<0.01; ***: p<0.001, significance of single compound vs Solvent is marked with § sign). (F) Western analysis of the co-controlled PPARα and ERRα target CPT1α from two corresponding liver samples with set-up as in (E). (G) Pie chart representation of the PPARα peaks (induced by GW7647) subdivided according to the proximity to an ERRα peak and top hit de novo motif enrichment on the genes, located in the promoter-TSS region, annotated with the >80% overlapping PPARα-ERRα peaks only (full list in Supplementary Fig. S5B).

### In silico genome-wide cistrome analysis identifies the ERRα motif underneath overlapping PPARα and ERRα binding sites within promoter regions

To find out whether PPARα target genes might be co-controlled by close-by binding of PPARα and ERRα, we performed overlays of our previously published PPARα cistrome dataset ^29^ with a publicly available ERRα cistrome dataset in murine hepatocytes ^30^, bearing in mind the limitation that only the first dataset entails a context of starvation. A total of 2532 binding sites for PPARα (after stimulation with GW) (Fig. 4G) and 9383 sites for ERRα (constitutively active) were retrieved. Almost half of those PPARα peaks (41.6%; 1054) had an ERRα binding site within 10 kb (Fig. 4G). Over one-fourth (294 out of 1054 peaks; 27.9%) of this segment showed an overlap of at least 1bp with an ERRα peak (Fig. 4G, Supplementary Table 2) and only a small proportion hereof overlapped more than 80% (49 peaks; 4.6%). The majority of the 294 PPARα-bound peaks that overlap with an ERRα peak were located in intronic and distal intergenic regions of the genome. Only a small fraction (16.7%) is retrieved in the promoter-TSS region (Supplementary Fig. S5A). Motif searches underneath overlapping peaks located in the promoter-TSS regions showed NR signatures with both ERR and PPARα motif enrichments (Supplementary Fig. S5A). Not unsurprisingly, a *de novo* motif search pointed to the short ERRα binding motif as the top hit for the maximally overlapping PPARα-ERRα peaks (Fig. 4G and Supplementary Fig. S5B). Upon assigning peaks to their closest gene, gene ontology analysis revealed enrichment of energy metabolism terms (Supplementary Fig. S5C). Despite using mouse liver cistrome datasets from different sources and experimental set-up, the analysis reveals that PPARα and ERRα peaks can overlap genome-widely.

### The context-dependent functioning of ERRα as a rheostat of PPARα-induced gene expression

To find out how combined ligand-responsive PPARα target genes are affected over time in a human cell context, we analyzed mRNA of HepG2 human hepatoma cells in a time course experiment. Both *CPT1*α and *PDK4* mRNA expression levels demonstrate a consistent gradual upregulation over time comparing GW/C29 to GW alone (Fig. 5A). The effect is notable from 24h onwards, in line with previous results (Fig. 3D) and most outspoken at 48h (left panel, cyan arrows). The mRNA of *UQCR10* and *Minos1* however follows a different pattern with overall lowered levels at 48h comparing GW/C29 to GW alone (orange arrows). Remarkably, in a non-hepatocyte cell model (L929sA, murine fibroblasts), we discovered a similar transcriptional regulation for a stably integrated PPARα-dependent luciferase reporter. This reporter contains the PDK4 enhancer region, verified to bind PPARα as previously revealed via ChIPseq ^29^. Here, crosstalk is apparent upon 24h of serum starvation (Fig. 5B), which is coincidentally the same context that also supported a stronger PPARα-ERRα interaction profile (Fig. 2A). Because of the evidence in literature for cholesterol as an endogenous ERRα agonist ^31^, we queried whether cholesterol addition could counteract the effects of an ERRα blockage by C29. Increasing amounts of cholesterol indeed compromised the ability of C29 to block PPARα-driven PDK4 enhancer-driven gene expression, in essence phenocopying the results obtained in the presence of serum. We verified that none of the inductions or treatments affected cell viability (Supplementary Fig. S6A, Fig. S6B).

**Fig. 5.**
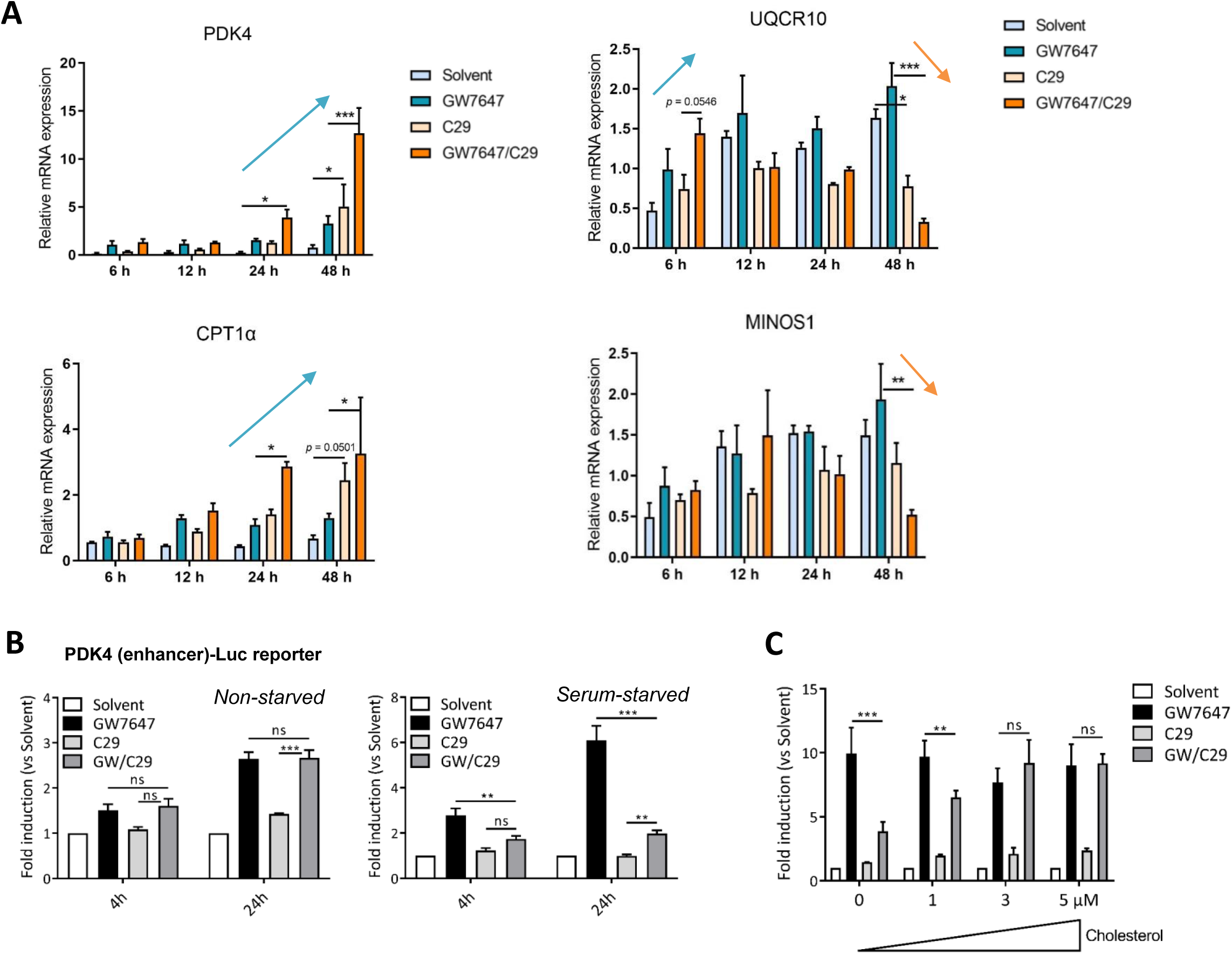
The cellular context and target gene identity co-determine ERRα’s ability to influence PPARα- mediated gene expression. (A) HepG2 cells were serum-starved overnight followed by treatment with GW7647 (0.5 µM) and/or C29 (5 µM) for 6, 12, 24 and 48h. qRT-PCR analysis shows relative mRNA expression levels of *CPT1a, PDK4, UQCR10* and *MINOS1,* using *GAPDH* and *TBP* as reference genes (Mean + SEM, n = 3). The significance analysis was performed by two-way ANOVA, followed by Tukey’s multiple comparisons test (*: p<0.05; **: p<0.01; ***: p<0.001). (B) L929sA cells with a stably integrated pPDK4-luc+, a PPRE-dependent promoter construct, were deprived from serum, or not, as indicated in the figure. After 4h, cells were stimulated with PPARα ligand GW7647 (GW, 0.5 µM) and/or ERRα inverse agonist C29 (5µM) for 4 or 24h. Promoter activities are expressed as relative induction factor versus Solvent (Mean + SEM, n=3). The significance of differences in reporter activity was evaluated with unbalanced two-way ANOVA, followed by multiple comparison using the Fisher’s LSD test (*: p<0.05; **: p<0.01; ***: p<0.001). (C) L929sA cells with a stably integrated pPDK4-luc+, a PPRE-dependent promoter construct, were, after 4h of serum-starvation with a cholesterol gradient as indicated, stimulated with PPARα ligand GW7647 (0.5 µM) and/or ERRα inverse agonist C29 (5µM) for 24h. Promoter activities are expressed as relative induction factor versus Solvent. Three independent replicates were performed (Mean + SEM). The significance of differences in reporter activity was evaluated with two-way ANOVA, followed by multiple comparison using the Fisher’s LSD test (*: p<0.05; **: p<0.01; ***: p<0.001).

### ERRα is recruited onto the chromatin by GW7647 near PPARα-driven target genes

Because we observed a consistent yet modest effect of transcriptional PPARα·ERRα crosstalk in 16h- fasted livers (Fig. 4E), we wondered whether a catabolic fasting state of 24h might enhance the crosstalk. As expected, blood metabolite analysis following 24h fasting showed high levels of ketone bodies, while lower trends of glucose levels were apparent in all fasted compared to fed conditions (Supplementary Fig. S7A). In contrast to the 16h fasting profile, however, gene expression analysis of livers upon prolonged starvation showed a lack of cooperativity comparing GW/C29 to GW for key metabolic pathway-controlling PPARα target genes (e.g. *Pdk4* and *Ehhadh*) (Supplementary Fig. S7B). This result may be indicative of a maximal gene expression plateau reached by the starvation-enhanced FA influx and/or GW treatment alone. We nevertheless pursued whether remnants of crosstalk between PPARα and ERRα, following single and combined ligand treatments, may be found at the level of DNA and went on to investigate the chromatin recruitment pattern of ERRα and/or PPARα. Western analysis showed weak PPARα protein signals in the ERRα ChIP (Fig. 6A), suggestive of PPARα’s presence in the pulled-down chromatin of fasted liver samples. Vice versa, even though poor anti-PPARα antibody quality did not allow us to build a chromatin recruitment profile for the same set of target genes, Western analysis of PPARα ChIP material did reveal ERRα protein. Based on our previous study using starved primary hepatocytes, we combined a view of PPARα cistromes with publicly available ERRα liver cistrome datasets (Supplementary Figure S8A). Despite that none of the latter datasets included starved conditions, the existence of overlaps of PPARα and ERRα peaks was suggested near PPARα-controlled promoter-transcription start site regions. Because we observed for *Pdk4* and *Ehhadh* trends indicative of combined ligand crosstalk in the fed state (Supplementary Fig. 7B, white bars), with levels for GW higher than for GW/C29, we expanded the analysis of DNA binding events also to the fed liver counterparts.

**Fig. 6.**
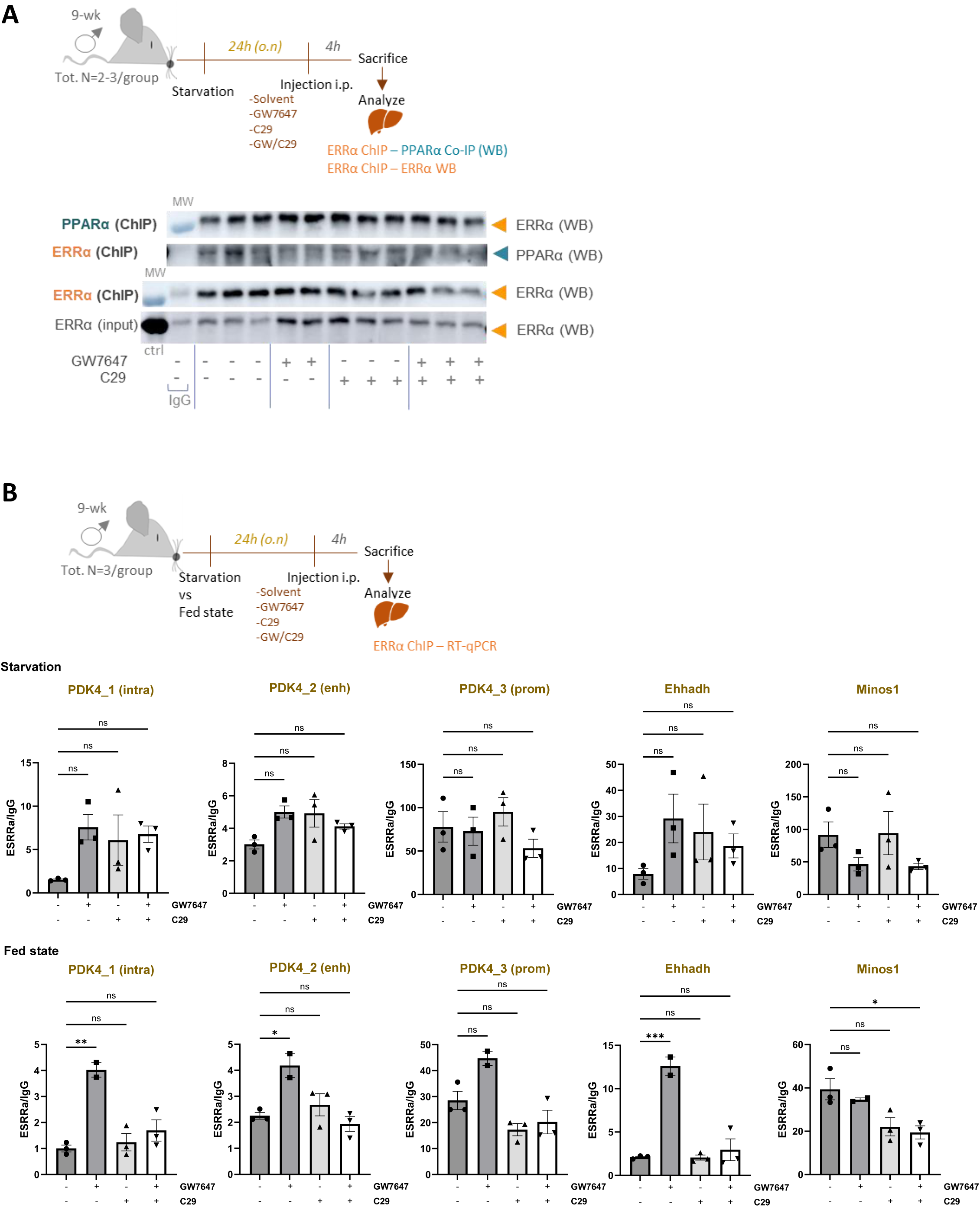
PPARα agonist treatment supports ERRα recruitment at the chromatin of PPARα-dependent promoters and enhancers. C57BL/6J mice (n=3), after 24h starvation, were treated for 4h with GW7647 (4mg/kg) and/or C29 (10 mg/kg) or were given solvent control, via i.p. injection. Collected livers were used for ChIP-Western analysis with anti-ERRα or anti-PPARα (A) and for ChIP-qPCR (B) which was performed with an anti-ERRα antibody versus IgG control. Results after immunoprecipitation were subtracted from the input and expressed as relative enrichment to the negative IgG control. The statistical significance was assessed using one-way ANOVA, followed by multiple comparison using the Tukey’s multiple comparison test (*: p<0.05; **: p<0.01; ***: p<0.001).

Following PPARα agonist treatment, we found significant ERRα recruitment at the chromatin of known PPARα-controlled promoters and enhancer DNA in fed livers (Fig. 6B, bottom panel and Supplementary Figure 8A for the different *Pdk4* primer localisations). Subsequent loss of ERRα recruitment upon combining GW7647 with the ERRα inhibitor C29 (Fig. 6B, bottom panel), indicative of a crosstalk mechanism at the level of DNA, may be caused by the inhibitor’s impact on ERRα levels (Fig. 6A) and/or a genuine loss of ERRα from (a complex on) the DNA. Exploration of ERRα binding in the prolonged fasted liver state showed, in line with the transcripts (Supplementary Fig. 7B), also less obvious ligand responses (Fig. 6B, upper panel). Still, overall amounts of recruited ERRα were at least equal or higher in a fasted liver state, for all PPARα responsive DNA regions tested (Supplementary Figure S8B). This result may suggest dominant effects of the catabolic fasted state over ligand effects. A summarizing model taking into account all findings throughout our study depicts how ERRα may function as a rheostat for PPARα dependent gene targets. The regulatory function of ERRα foremost depends on the target itself and is next influenced by fasted versus fed liver states, whereby loss of ERRα recruitment upon ERRα inhibition was surprisingly most apparent in the fed state (Fig. 7).

**Fig. 7.**
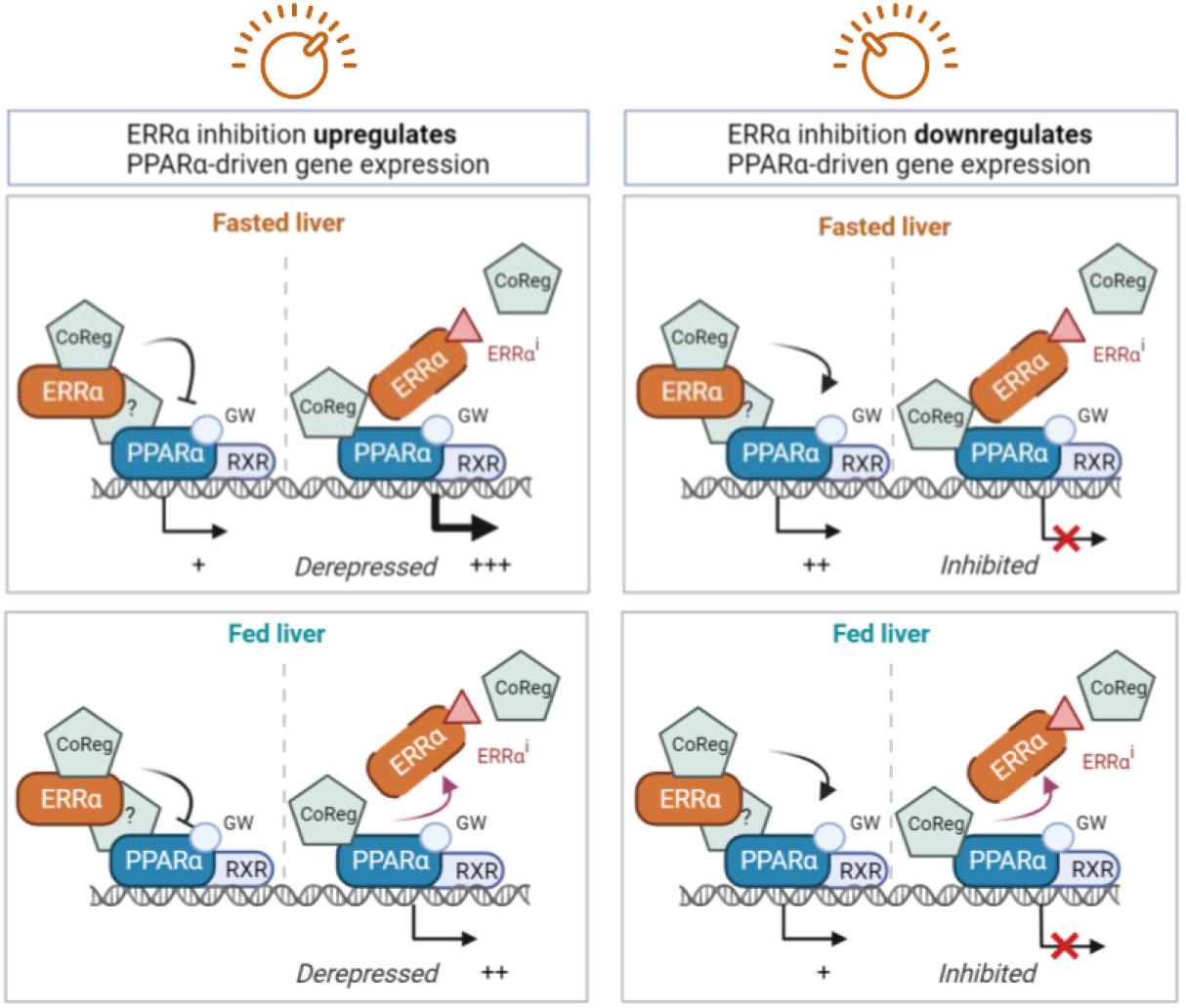
Model for opposite ERRα regulation of PPARα-dependent target genes in a context-dependent manner. In the fasted liver state (top panels), both constitutively active ERRα and C29-inhibited ERRα may remain in a complex together with GW7647(GW)-liganded PPARα-RXRα to finetune the expression of different PPARα targets. The question mark points to PGC1 coregulators as likely bridging factors. The left panel exemplifies a transcriptional regulation as seen for PDK4 promoter activity whereas the right panel may reflect a regulation as observed for the isolated PDK4 enhancer. In a fed liver state, patterns of transcript regulation were fairly similar to the fasted state, save a clear loss of ERRα recruitment from the chromatin following ERRα inhibition. The model was created via biorender.com.

## Discussion

The data presented here brings forward several lines of evidence that a PPARα·ERRα interaction, likely bridged by a coactivator such as PGC1α and inclusive of RXRα, represents a functional transcriptional complex able to influence PPARα-driven gene expression. First, ERRα was picked up as one of the top hits in a MAPPIT mammalian two-hybrid screen designed to capture not only strong interactions but also weaker and transient ones (Fig. 1A). While cell-based experiments consistently showed a strong interaction (Fig. 1B-C, Fig. 2B), likely nuclear (Fig. 1D-E, Fig. 2C, Supplementary Fig.S1B) and involving the LBD of PPARα (Supplementary Fig. S2B), only a weak direct PPARαLBD·ERRα interaction emerged using *in vitro* binding assays (Fig. 1H, Supplementary Fig. S1D). In sharp contrast to GST-pulldown assays (Fig. 1F, Supplementary Fig. S1C), the functional interaction within cells revealed a clear PPARα ligand dependency (Fig. 1A-C, Fig. 2A-B, Supplementary Fig. S2B). Random mutagenesis interaction studies (Supplementary Fig. S2D-G) allowed detailed mapping of the binding interface of PPARα and revealed a discrete patch encompassing its classic coactivator binding site (Fig. 2D). Subsequent modeling suggested the interaction involved the C-terminal AF2 domain of ERRα (Supplementary Fig. S2G), which was confirmed by a directed mutagenesis strategy (Fig. 2F). A role for PGC1 family members was suspected, given these coregulators are well-characterized protein ligands of ERRα ^10, 22, 26, 32^, and known to be upregulated under various physiological cell stressors e.g. a fasted liver ^33, 34^. Our data further align with a previous study showing that ERRα binding to PGC1α requires the C-terminal AF2 domain of ERRα ^33^. In support of a functional interplay involving PPARα, coregulator profiling showed that the loss of PGC1 coactivators to ERRα upon PPARα ligand activation coincided with a gain of these coregulators to PPARα (Fig. 3B).

Second, ERRα coactivated ligand-activated PPARα in heterologous Gal4-dependent promoter-reporter assays (Fig. 2C). This activity could be suppressed by either using pharmacological inhibitors of ERRα or using the AF2 MLM mutant of ERRα (Fig. 2G). Inverse agonists of ERRα were previously shown to disrupt the interaction between ERRα and PGC1 coregulators by causing helix 12 of ERRα to shift into its own coactivator groove ^13, 35^, and the latter interaction was shown to also rely on the AF2 of ERRα^26^. In line, *in vitro* assaying showed that both inverse agonists used in the study lowered the already weak PPARα·ERRα interaction (Fig. 1H).

The specific set-up of MAPPIT, which retrieves protein-protein interactions in the cytoplasmic compartment where coregulator concentrations are expectedly lower than in the nucleus, may have given an advantage to enrich for a transient, direct interaction that may still be disfavored in the nucleus. Importantly, overexpression of PGC1α WT or single L2 and L3 mutants hereof all show specific enhancement of the MAPPIT-captured PPARα·ERRα interaction, except for the L2A/L3A double mutant which can no longer bind to both PPARα or ERRα ^26^ (Fig 3A). Our results thus suggest that an intact L3 region described to exclusively interact with ERRα ^26^ is sufficient to support the PPARα·ERRα interaction. How come a basal PPARα·ERRα interaction profile can still be observed in the presence of the L2A/L3A double mutant, may be explained either by residual direct interaction or by the presence of an endogenously bridging PGC1α, or even other coregulators that can shuttle between cytoplasm and nucleus. Collectively, the interaction data so far align with the involvement of coregulators to strengthen *in cellulo* an otherwise unstable, weak, or transient interaction. Another striking feature of the PPARα·ERRα interaction was that it could be enhanced by depriving the cell culture of serum (Fig. 2A). Under the same conditions, scrutiny of a reporter gene composed of the PPARα ligand-responsive PDK4 enhancer coupled with luciferase showed that inhibition of ERRα activity compromised enhancer activity, attributing a co-activating role for ERRα in this particular context (Fig. 5B). Adding increasing amounts of cholesterol to the cell culture completely restored liganded PPARα’s full transcriptional capacity, even in the presence of the ERRα inhibitor (Fig. 5C). These findings suggest that serum deprivation and more specifically the lack of cholesterol in the cells may drive a functional interaction between ERRα and activated PPARα on the PDK4 enhancer. As such, our data would agree with a previous report identifying cholesterol as an endogenous ERRα ligand ^31^ and also with a study wherein cholesterol was suggested to stabilize the interaction between PGC1α and helix 12 of ERRα ^36^. Because serum starvation can modulate various kinase/phosphatase-controlled signaling pathways ^37, 38^ and given a role for posttranslational modifications has been reported both for PPARα and ERRα ^2, 39^, the precise molecular basis explaining an enhanced PPARα·ERRα interaction and activity profile following serum starvation awaits further research.

Third, a role for PPARα·ERRα in regulating hepatocyte gene regulatory processes is suggested by differential mRNA expression profiles of PPARα-driven target genes in the absence and presence of pharmacological ERRα inhibitors (Fig. 3D, Fig. 4A-E). Following RNA sequencing of starved murine livers, interaction term analysis unsurprisingly connected PPARα·ERRα crosstalk to mitochondrial FAO and oxidative phosphorylation pathways (Fig. 4B-C). ERRα’s functioning as a (mild) repressor of PPARα-driven *PDK4* gene expression in starving hepatocytes seemingly contrasts with an activating role of ERRα in non-hepatocyte models and isolated *PDK4* promoter studies ^11, 12^. Opposite roles of ERRα were however previously identified in a context of liver gene expression, where ERRα and PGC1α costimulatory effects on mitochondrial gene expression coincided with a role for ERRα as a transcriptional repressor of the PEPCK gene^5^. Pharmacological inhibition of ERRα, reported to destabilize and decrease hepatocyte ERRα protein levels ^40^, enhanced mRNA levels of GW-induced prototypical PPARα-driven target genes including *PDK4* (Fig. 4E). This enhancement did not correlate with higher PPARα transcript or protein levels following ERRα inhibition (Fig. 4A, Fig. 4F). Increased CPT1α protein levels were similarly found when combining C29/GW, and coincided with lower ERRα protein levels (Fig. 4F). In this context, the data are suggestive of a role for ERRα as a transcriptional corepressor of PPARα functioning, whereby the brakes are relieved following the inverse agonist’s actions on ERRα. The latter event can moreover operate at multiple non-exclusive levels, via loss of direct or indirect interactions with PPARα and/or a bridging coregulator, and/or by loss of ERRα protein. Time-course experiments of some PPARα-driven genes that emerged from the RNAseq analysis, showed a shift in mRNA regulation profiles over time (Fig. 5A). These profiles allow for many secondary effects and thus reflect additional layers of complex regulations, for instance by autoregulatory loops controlling both PPARα and ERRα ^41, 42^. Hence, under the circumstances wherein ERRα behaves as a corepressor, loss of ERRα functionality and gain of PPARα protein function would be expected to spur downstream PPARα target gene expression, as may be the case for both PDK4 and CPT1α. Intracellular crosstalk mechanisms between PPARα and ERRα are relevant also in other systems a.o. in kidney cells and (cardiac) myocytes, where a transcriptional influence of each other’s gene expression and a DNA binding-dependent suppression of the cardiac ERRα transcriptome by PPARα was observed ^10, 43^ ^44^.

A final piece of data in support of the physiological relevance of a PPARα·ERRα crosstalking complex in liver was the retrieval of ERRα at the chromatin of ligand-activated PPARα-controlled target genes (Fig. 6A-B). Chromatin-immunoprecipitated ERRα was present in a PPARα ligand-dependent manner at various promoter and enhancer regions of select PPARα-controlled targets. PPARα ligand-dependent ERRα recruitment was significant in fed livers while a trend was observed also in starved livers (Fig. 6B), pointing to a conserved layer of control. Taken together, our data support PPARα·ERRα interactions between both endogenous transcription factors, direct or indirect, with both receptors likely present within the same transcriptional complexes, both in fasted and fed liver states, yet with the latter state more susceptible to ERRα inhibition (Fig. 6B). It remains to be investigated how genome-wide binding profiles of other involved factors may vary comparing GW vs GW/C29, for example RXRα as an established partner protein of PPARα (Fig. 1G) as well as PGC1α as a cofactor for both receptors, in livers under starved versus fed conditions.

In sum, ERRα manifests as a transcriptional modulator of liver PPARα functioning, a so-called “rheostat”, capable of either dampening or activating PPARα-driven gene expression depending on the target and serving to finetune the expression of PPARα-driven genes involved in metabolic gene expressions in the liver. Multiple layers of crosstalk mechanisms between PPARα and ERRα may coexist as a mechanism hepatocytes use to adapt to changes in nutrient availability. The PPARα-ERRα axis is therefore a likely system prone to become imbalanced in a diseased liver state, such as e.g. metabolic dysfunction-associated fatty liver disease (MAFLD, formerly NAFLD)^45^. Indeed, loss of ERRα function protected from high fat diet-induced body weight gain and MAFLD ^46^. In further support, a recent study on hepatocyte loss of the E3 ligase FBXW7 showed detrimental downstream consequences for metabolic transcriptional axes jointly controlled by PPARα and ERRα ^47^, suggestive of a solid basis for co-targeting strategies of both receptors in MAFLD. The understanding of NR-centered crosstalk mechanisms is becoming highly relevant, especially because polypharmacological approaches targeting multiple metabolic NRs are increasingly considered to combat various metabolic disorders.

## Materials and methods

### Compounds & reagents

PPARα ligands GW7647 (GW) and pemafibrate (pema) were purchased from Sigma-Aldrich and Bioconnect, respectively. A stock solution was prepared in DMSO (5mM) and stored at −20°C. XCT790 was purchased from Sigma-Aldrich (cat. Nr. X4753). Compound 29 (C29), the inverse agonist of ERRα, was synthesized as described ^13^(patent US2006014812A1). Stock solutions of XCT790 or C29 were prepared in DMSO (10mM) and stored at −20°C protected from light.

### Cell Culture

HEK293T and L929sA cells were cultured in DMEM medium containing 10% FBS. HepG2 cells were cultured in DMEM or Opti-MEM containing 10% FBS or 10% DCC-FBS. Primary hepatocytes were isolated from C57BI/6J WT male mice as described in ^29^, and seeded on iBidi µSlides (precoated with Collagen type I) in Williams medium (+ additives, see ^29^). 2h after attachment, medium was replaced (without additives) and left for another 2h before stimulation.

### Animals

All experiments were approved by the institutional ethics committees for animal welfare of the Faculty of Sciences and Faculty of medicine and Health Sciences, Ghent University, Belgium (Ethical dossier numbers ECD 14/83 and ECD 14/84). Male C57BL/6J mice were purchased from Janvier (Le Genest-St. Isle, France). Mice were housed in a temperature-controlled, specific pathogen free (SPF) air-conditioned animal house with 14 and 10h light/dark cycles and received food and water ad libitum. All mice were used at the age of 9-11 weeks. During starvation experiments, food was taken away either in the morning or afternoon for time periods to match a 24h or 16h starvation. Between 9-10am the next day, mice were injected intraperitoneally according to body weight with solvent control (Ringer solution), GW7647 (4mg/kg) and/or Compound 29 (10mg/kg). GW7647 was prepared as a solution of 8 mg/ml in DMSO. Similar, C29 was dissolved in DMSO at a concentration of 20 mg/ml. GW7647 and C29 were further diluted to 4 mg/kg and 10 mg/kg, respectively, in Ringer solution. 4h after injection, mice were euthanized via cervical dislocation. Liver samples were isolated and stored in RNAlater (Ambion) at −20°C.

#### Biochemical analysis

Blood glucose, lactate and ketone body levels were measured in tail blood with the use of OneTouch Verio glucose meter (LifeScan), Lactate Plus meter (Nova Biomedical), and Freestyle Precision Neo meter (Abbott), respectively.

### Receptor protein expression and purification

*ERRa -* A plasmid encoding FL-ERRα was ordered from GenScript. The full-length human ERRα (residue 1-423) gene with NCoI and NotI restriction sites was cloned into a pET28b(+) vector to include a C-terminal His-tag. Transformation of E. Coli BL21 (DE3) competent cells with the plasmid was performed with heat-shock. A single colony was picked and transferred to 25 mL LB medium supplied with 50 µg/mL kanamycin. This culture was incubated overnight in a shaking incubator at 37°C. The small cultures were transferred to 2 L Terrific Broth (TB) medium supplied with 0.05% antifoam SE-15 (Sigma Aldrich) and 50 μg/ml kanamycin. Using 0.5 mM IPTG, protein expression was induced when an OD_600_ of 0.8 was reached. Protein expression proceeded overnight at 18°C at 150 rpm. After 15 hours, the cell suspension was centrifuged at 10.000 RCF for 10 minutes at 4°C. The resulting cell pellet was resuspended in lysis buffer (20 mM Tris (pH=7.9), 500 mM NaCl, 10 mM imidazole, 25 U/ml Bezonase® Nuclease (Millipore) and one cOmplete™ Protease Inhibitor Cocktail tablet (Roche) per 25 ml cell suspension). An Emulsiflex-C3 homogenizer (Avestin) was used to lyse the cells and the lysate was cleared using centrifugation at 40.000 RCF for 40 minutes at 4°C. The supernatant was loaded on a 1 ml Ni-NTA Superflow cartridge (QIAGEN). The column was washed with 10 column volumes (CVs) of buffer A (20 mM Tris (pH=7.9), 500 mM NaCl and 10 mM imidazole) and 10 CVs of buffer A supplied with 45 mM imidazole. The purified protein was eluted using 8 CVs of buffer A with 200 mM imidazole. This fraction was then dialyzed overnight to a buffer containing 50 mM Tris (pH=7.9), 100 mM NaCl, 50 µM EDTA and 20% glycerol. Subsequently, the solution was concentrated using an Amicon® Ultra centrifugal filter with a 10 kDa cutoff (Millipore). The product was aliquoted, flash-frozen and stored at −80°C. The purity of the product was assessed using SDS-PAGE analysis.

PPARα-A plasmid encoding the STREP-PPARα LBD was ordered from GenScript. A sequence encoding for the human PPARα LBD (residue 200-468) with an N-terminal Strep-tag®II was cloned into a pET15b vector using NdeI and XhoI restriction sites. E. Coli BL21 (DE3) competent cells were transformed with the plasmid using heat-shock. A single colony was used to start a culture of 25 ml LB medium supplied with 100 µg/ml ampicillin, which was incubated overnight at 37°C. The starter cultures were transferred to 2 L of TB medium supplied with 0.05% antifoam SE-15 (Sigma Aldrich) and 100 µg/ml ampicillin. Protein expression was induced using 0.5 M IPTG when an OD_600_ of 0.8 was reached. Expression continued overnight at 15°C and 150 rpm. The cell pellet was collected by centrifugation at 10.000 RCF for 10 minutes at 4°C and resuspended in lysis buffer (50 mM Tris (pH=7.8), 300 mM NaCl, 20 mM imidazole, 10% glycerol, 25 U/ml Bezonase® Nuclease (Millipore) and one cOmplete™ Protease Inhibitor Cocktail tablet (Roche) per 25 ml cell suspension). Cells were lysed using an Emulsiflex-C3 homogenizer (Avestin) and the lysate was centrifuged at 40.000 RCF for 40 minutes at 4°C. A 5 ml Strep-Tactin®XT Superflow® high capacity cartridge was equilibrated with buffer B (50 mM Tris (pH=7.8), 300 mM NaCl, 20 mM imidazole, 10% glycerol). The cleared solution was loaded onto the column, which was subsequently washed with 10 CVs of buffer B. The purified protein was eluted using 5 CVs of buffer B supplied with 50 mM EDTA. The sample was dialyzed overnight to buffer C (20 mM Tris (pH=7.8), 150 mM NaCl, 5 mM DTT and 10% glycerol). The dialyzed solution was loaded on a Superdex 75 pg 16/60 size-exclusion column (GE Life Sciences) using buffer C as a running buffer. The elution fractions were analyzed using high-resolution mass spectrometry (Xevo G2 Quadrupole Time of Flight) and SDS-PAGE. Fractions containing the correct mass were combined, concentrated and stored at −80°C.

### GST-pulldown

The pulldown was performed with GST-PPARα, GST-PGC1α (positive control, containing the 1-293 fragment of the full-length protein; kindly gifted by Dr. A. Kralli, Johns Hopkins University, Baltimore), GST-5HT7 or GST-empty (neg. ctrls) according to the protocol described in ^29^. *In vitro* transcribed and translated FLAG-tagged ERRα protein was made using the TnT reticulocyte reaction (Promega cat #L1170, 2h on 37°C) or the TnT wheat germ extract (cat #L5030, 90’ on 30°C), and added (20µl/sample) to the beads solution, together with 5µM GW or solvent control (DMSO). Immunostaining was performed using the anti-FLAG M2 (F3165, Sigma Aldrich) at 1:1000 and anti-GST antibody (ab9085, Abcam) at 1:500.

### In vitro His-tag pulldown

To remove traces of EDTA and to ensure proper folding of FL-ERRα, the protein samples of FL-ERRα and the PPARα LBD were both buffer exchanged to buffer D (20mM Tris (pH=8.0), 300mM NaCl, 15mM imidazole, 100 µM ZnCl_2_) using PD SpinTrap G-25 columns (GE Healthcare). Ni-NTA magnetic agarose beads (QIAGEN) were added to His-tagged FL-ERRα and His-NanoLuc (= neg. ctrl), and incubated for 1h at 4°C. The tubes were placed on a DynaMag-2 magnetic separator (Thermo Fisher) for 1’ to remove the solution. Beads were washed with an excess of buffer D before being placed back on the magnetic separator and extracting the solution. Next, the PPARα solution was added to the magnetic beads, following 1h incubation at 4°C. The solution was removed using the magnetic separator and the wash step was repeated. Finally, buffer D supplied with 200mM imidazole was used to elute the product. SDS-PAGE was used to analyze sample composition.

### Proximity ligation assay (PLA)

PLA was performed using the DuoLink In Situ Red Starter Kit Mouse/Rabbit (DUO92101, Sigma). µ-slide 8-wells (Ibidi) seeded with HepG2 cells were fixed, permeabilized and blocked with the Duolink Blocking Solution for 1h at 37°C. Next, the slides were incubated overnight at 4°C with mouse anti-PPARα (1µg/ml, sc398394, Santa Cruz) and rabbit anti-ERRα (1µg/ml, ab76228, Abcam) in Duolink Antibody Diluent, followed by 1h incubation at 37°C with Duolink In Situ PLA Probe anti-rabbit PLUS and anti-mouse MINUS (10x dilution). All washing, ligation and amplification steps were performed following the manufacturer’s instructions.

### Reporter assays

L929sA cells were stably transfected (by standard CaPO_4_ procedure) with p(PDK4)-Luc+ reporter gene construct. This construct was generated by amplification of the PPRE peak sequences in the PDK4 promoter region from mouse liver genomic DNA and ligation in a pGL3-basic vector. 4h before induction, medium was replaced by fresh medium in the presence or absence of serum, as indicated in the Figure legends. For the Gal4 experiments, cells grown in DMEM/FCS were transfected with the Gal4-reporter plasmid, Gal4(DBD)-control, Gal4(DBD)-PPARα or Gal4(DBD)-PPARαLBD chimera, and ERRα plasmids. After 4h in the presence of Opti-MEM with 10% DCC-FBS, cells were induced as indicated, followed by luciferase assays according to the protocol of Promega Corp. For each biological replicate, luciferase measurements were performed at least in triplicate and normalized, where possible, by measurement of β-galactosidase (β-gal) levels with the Galacto-Light kit (Tropix). Light emission was measured with the TopCount NXT luminometer or EnVision (Perkin-Elmer).

### qRT-PCR

HepG2 cells were induced as indicated and total solvent concentration was kept similar in all conditions. The cells were serum-starved 4h before induction. RNA was isolated using RNeasy Micro Kit (Qiagen). RNA from the murine livers was isolated using TRIzol Reagent, according to the manufacturer’s protocol (Invitrogen). Next, mRNA was reverse transcribed to cDNA with the PrimeScript RT kit (TaKaRa). cDNA was analysed by real-time PCR with the Light Cycler 480 SYBR Green I Master Mix (Roche). A set of 3 household genes has been used to normalize (Cyclophilin, β-actin and HPRT1). Primer sequences are included in a Supplementary Table 3.

### SDS-PAGE and Western Blot analysis

After washing with ice-cold PBS, cell lysates were prepared using SDS sample buffer followed by Western Blotting and antibody probing procedures according to the guidelines of the company for the respective antibodies (anti-PPARα: H-2, sc-398394, 1:1000, Santa Cruz Biotechnology; anti-CPT1α: ab128568, 1:1000, Abcam; anti-ERRα: ab137489, 1:1000, Abcam). Imaging was done using KODAK films or the Amersham Imager 680 equipment.

### Co-immunoprecipitation

HEK293T cells, transiently transfected (CaPO_4_) with the plasmids as indicated, were serum-starved overnight, followed by 3h stimulation. Next, cells were lysed (50mM Tris-HCl pH7.5, 125mM NaCl, 5% glycerol, 0.2% NP40, 1.5mM MgCl_2_ and Complete Protease Inhibitor Cocktail (Roche)) and incubated overnight with anti-FLAG beads (anti-FLAG M2 affinity gel, Sigma Aldrich) on a rotor at 4°C. These beads were blocked beforehand for 1h at 4°C, using undiluted StartingBlock^TM^ (TBS) Blocking buffer (Thermo Scientific). After washing, the samples were eluted using Laemmli buffer, boiled for 5’ at 95°C and stored at −20°C. Finally, Western Blotting and antibody probing procedures were performed according to the guidelines of the company for the respective antibodies anti-HA (Roche), anti-FLAG (Sigma Aldrich), anti-E (Phadia) and anti-actin (as loading control) antibody (Sigma Aldrich). Imaging was done using KODAK films or the Amersham Imager 680 equipment.

### Immunofluorescence - Image capture and analysis

HepG2 cells, seeded on poly-L-lysine-coated μ-slides (Ibidi) and serum-deprived overnight, were induced as indicated. After fixation, endogenous PPARα was visualized using mouse anti-PPARα (H-2, sc-398394) antibody (Santa Cruz Biotechnology), followed by Alexa Fluor 568 anti-mouse IgG (Molecular Probes, Invitrogen). Endogenous ERRα was visualized using rabbit anti-ERRα (ab137489) antibody (Abcam), followed by Alexa Fluor 488 anti-rabbit IgG (Molecular Probes, Invitrogen). Cell nuclei were stained with DAPI DNA staining (300nM, Invitrogen).

Confocal images (8-bit) were captured with an LSM880 confocal microscope equipped with an Airyscan detector (Zeiss, Jena, Germany). Images were taken in super-resolution, FAST mode by using a Plan-Apochromat 63x/1.4 oil objective (frame size: 2544 × 2544 pixels, pixel size: 35 nm × 35 nm). AF 488 was excited using the 488 nm line of an Ar laser (5%) and emission was captured between 495 and 550 nm. AF 568 was excited by a diode laser at 561 nm, and emission was detected using a combined filter (BP570-620+LP645). To study nuclear colocalizations, Z-sections were made every 159 nm. Images were calculated through pixel reassignment and Wiener filtering by using the built-in “Airyscan Processing” command in the Zen software. Thresholded Pearson Correlation Coefficients (PCC) were calculated after segmentation in 3D using Volocity software (Quorum Technologies). PPARα positive objects (groups of pixels with intensity values above a pre-defined threshold and larger than 4 pixels) were segmented out and PCCs were calculated for these objects. The average PCC value over all objects in a particular image stack was calculated. For each condition, 6 image stacks were analyzed (each image stack contained on average 8 cells).

### Array and binary MAPPIT

Mammalian protein-protein interaction trap (MAPPIT) is a two-hybrid system based on the restoration of a dysfunctional cytokine receptor signaling pathway. The principle of a conventional MAPPIT is depicted in Fig. 1b. The bait protein is C-terminally fused to a mutant leptin receptor, of which three conserved tyrosine (Y) residues are mutated to phenylalanine (F). Binding of leptin activates the receptor-associated Janus kinases (JAK). However, due to the Y to F mutations, the receptor is unable to recruit and activate signal transducers and activators of transcription (STAT)3 proteins and induce reporter activity. The prey protein is coupled to the C-terminus of glycoprotein 130 (gp130), a receptor fragment that contains four functional STAT3 recruitment sites. Upon bait-prey interaction, the JAK/STAT signaling pathway is restored and leads to STAT3-dependent reporter activity. Array MAPPIT and the preparation of the prey and reporter reverse transfection mixture was previously described^16^. 8.569 full-length human ORF preys, selected from the human ORFeome collection version V5.1 (http://horfdb.dfci.harvard. edu/hv5), constituted the screened prey collection. The binary MAPPIT analysis was performed as described earlier ^29^. In case of serum starvation, medium was replaced by DMEM without FBS at the time of stimulation. The generation of the PPARα-bait plasmid (pCLG-PPARα), the negative control bait (pCLG-eDHFR), the empty prey control (pMG1) and the pXP2d2-rPAP1-luciferase reporter have been described previously ^29^. The ERRα prey plasmid (pMG1-ERRα) was created by Gateway transfer of the full-size ERRα ORF, obtained as entry clone from the hORFeome collection (hORFeome version V8.1), into the Gateway compatible MAPPIT prey destination vector pMG1 as described earlier ^16^. The PPARα-LBD bait plasmid (pCLG-PPARα-LBD) was generated by substituting FL-PPARα-encoding sequence of the pCLG-PPARα vector with the LBD coding sequence of PPARα. The triple mutated ERRαMLM (M417A, L418A and M421A) was made using site-directed mutagenesis on the pMG1-ERRα plasmid.

### Random mutagenesis – MAPPIT

The PPARα-LBD sequence in the bait pCLG-PPARα construct was randomly mutated by error prone PCR, after which single mutants were selected and analysed in 384-well format MAPPIT assays as previously described ^48^. HEK293T cells were transfected (CaPO_4_) with ERRα prey plasmid, the different mutant PPARα-bait plasmids, and the pXP2d2-rPAP1-luciferase reporter. After 24h, half of the wells were stimulated with leptin (100ng/ml) in combination with GW (0.5µM), the other half only with GW. Cells were lysed and luminescence was measured using the EnVision plate reader (Perkin-Elmer). Comparison of the luminescence signal of the mutants with the wild-type PPARα bait resulted in the relative MAPPIT signal. The threshold for mutants that break the interaction (under 25% of wild-type interaction), and stimulating mutants (above 150%) were set based on the distribution of the wild-type PPARα-ERRα interactions. The binding effect of the mutations was mapped and visualised on the PPARα-LBD crystal structure using the UCSF Chimera package (University of California, San Francisco). More precisely, the mutations were mapped on the agonist (GW590735)-bound PPARα-LBD in complex with a peptide derived from the coactivator SRC-1 (PDB: 2P54), or on the antagonist (GW6471)-bound PPARα-LBD in complex with a peptide derived from the corepressor SMRT (PDB: 1KKQ). The effect of the mutation on the protein stability was calculated using FoldX with X-ray structure (PDB: 3VI8) of PPARα-LBD as template.

### Interaction model

In the apo-PPARγ-LBD structure (PDB: 1PRG), helix 12 of a first LBD binds in the coactivator pocket of a second LBD ^49^. The model for the PPARα-ERRα interaction (Fig. 2f) was obtained, using Yasara structure, by superposing the PPARα-LBD structure (PDB: 2P54) on the second LBD in the apo-PPARγ-LBD structure. The ERRα-LBD structure (PDB: 3D24) was superposed on the first LBD in the apo-PPARγ-LBD structure. Its helix 12 was separately superposed on helix 12 of the same PPARγ-LBD. The loop between helix 12 and the rest of the ERRα-LBD was energy minimized, followed by energy minimization of the entire PPARα-LBD - ERRα-LBD complex.

### RNA sequencing data analysis

RNA sequencing libraries were generated in biological triplicate using the TruSeq stranded mRNA protocol. Libraries were subjected to single-end 100bp sequencing on Illumina NovaSeq6000, yielding 12-20 million reads per sample. Subsequent data analysis was performed using a dedicated Snakemake pipeline. Briefly, the sequencing reads were quality controlled with FastQC (version 0.11.9). Next, Trim-Galore (version 0.6.6-0) was used to trim low-quality ends from reads (with phred score < 30), in addition to adapter removal. Following another quality control of the trimmed data, reads were pre-mapped to PhiX genome using STAR (version 2.7.6a) with a genome index setting -- genomeSAindexNbases 5, disabled splicing and a maximum of 2 mismatches allowed. The PhiX-unmapped reads were then aligned to the mouse genome GRCm38 using the splice junctions of the Ensembl 101 annotation, allowing only uniquely mapped reads and a maximum of 4 mismatches. The position-sorted output BAM files were converted to count data using HTSeq (version 0.12.4) in the ‘union’ mode. Every sample was present in technical replicates distributed over several Illumina flowcell lanes. First, outliers in the replicate count data were evaluated using R package ggbiplot (version 0.55), allowing analysis to proceed minus 1 DMSO outlier. Finally, read counts were summed per gene and per sample using reshape2 (version 1.4.4), tidyr (version 1.1.2) and dplyr (version 1.0.2). Differential expression analysis was conducted using DESeq2 R package (version 1.28.1). Genes with less than 50 reads in all replicates of at least one condition were removed prior to analysis. DESeq2 automatic independent filtering was enabled, therefore genes having a low mean normalized count were further filtered out and marked by an adjusted p-value set to NA. Pairwise contrasts of interest between differentially treated samples were retrieved at a significance level alpha 0.05, corresponding to Wald-test adjusted p-value (FDR) cutoff. Subsequently, an interaction term was added to the design formula to identify gene expression signatures attributed to the combined treatment with GW and C29. Log2 fold changes or, alternatively, normalized counts were compared, clustered and presented as heatmaps using the pheatmap package (version 1.0.12). Gene set enrichment analysis (GSEA) of gene ontology (GO) terms and KEGG categories, as well as GO enrichment analysis (when applicable to unsorted gene lists), were performed using clusterProfiler (version 3.16.1) according to mouse gene annotation package org.Mm.eg.db (version 3.11.4) with an adjusted p-value cutoff of 0.05. Enriched terms and pathways were further visualized using enrichplot (version 1.8.1) and pathview (version 1.28.1). Venn diagrams were created using eulerr (version 6.1.0). Other visualizations were generated using ggplot2 (version 3.3.2), ggrepel (0.8.2), ggsci (2.9) and RColorBrewer(1.1-2).

### ChIP-seq data analysis

Computational analysis was performed on raw read data from published ChIP-seq data, available in the short read archive (SRA) (PPARα: run SRR2043161 (control: run SRR2043176), ERRα: SRR650763 (control: SRR650764)). Reads were mapped to the mouse reference genome (mm10) using the Bowtie2 aligner (version 2.3.5) with settings “-t -p 4 –S”. Peak calling was performed using MACS (version 1.4.2) using matched input sample controls and requiring a p-value ≤ 1e-8 (with parameters: -g mm -p 1e-8 --bw 150 -B -S). The BEDOPS software (BEDOPS --element-of) was used to determine PPARα and ERRα peaks that were in close proximity or overlapping. To find peaks within a distance of 10 kb the ‘--range’ was used to extend the regions specified in the BED files. Overlapping peaks were determined with --element-of 1 or --element-of 80%. Peak-gene annotation, meaning assignment to the closest gene and classification according to genomic location, was performed using HOMER. Likewise, HOMER was also applied to perform motif finding using default settings. The genes that were annotated with peaks found to be located in a promoter-TSS regions were subjected to gene ontology and KEGG pathway enrichment using Enrichr.

### Statistics

Statistical analyses were carried out using GenStat v18 and GraphPad Prism v7 Software. Analysis of MAPPIT data was performed using one- or two-way ANOVA with REPLICATE as blocking factor, as implemented in Genstat. In case of missing values, the unbalanced design was applied. Post-hoc analysis using T statistics was performed assessing the significance of pairwise comparisons. A HGLMM (fixed model: poisson distribution, log link; random model: gamma distribution, log link) as implemented in Genstat has been fitted to the qPCR data of multiple genes jointly. The linear predictor vector of the values can be written as follows: log(*μ*) = *η* = **X***β* + **Z**ν, where the matrix X is the design matrix for the fixed terms (GENE, INDUCTION) and their interaction, *β* is their vector of regression coefficients, **Z** is the design matrix for the random term, and ν is the corresponding vector of random effects (INDIVIDUAL). T statistics were used to assess the significance of effects (on the transformed scale) by pairwise comparisons to the reference level (as indicated in the Figure legends). Estimated mean values and standard errors (SE’s) were obtained as predictions from the HGLMM, formed on the scale of the response variable.

## Data Availability

The RNAseq dataset generated during the current study is available in the GEO repository with the following accession GSE200658, accompanied by a reviewer access token “qfezeqmeffyjjqn”.

### Author contributions

KDB and SJD designed the study; SJD, JTh, TV, YL, RDV, LDC, DR, EV, and EVH performed experiments; DF analysed the RNA sequencing data; ST performed the ChIP-seq analysis; EVH assisted with the IF set-up and analysis; LDC and TV performed and analysed *in vivo* experiments; FP contributed with the random mutagenesis analysis, interaction data analysis; LB contributed with structural insights; SJD, JTh, TV, YL, EV, LDC, DR, DF, ST, LB, FP, BS, CL, JTa and KDB analysed and/or interpreted data; SJD and KDB drafted the manuscript; DF, BS, LB, CL and JTa discussed/advised on data and helped revising the manuscript. All authors approved the final version.

**Supplementary information** accompanies this paper

## Acknowledgements

SJD was supported by a fellowship of Fonds voor Wetenschappelijk Onderzoek (FWO)-Vlaanderen and Bijzonder Onderzoeksfonds (BOF) at UGent. Research at the labs of KDB and LB was funded by an FWO research project grant G015321N. EV was supported by a UGent-BOF grant. Research in the lab of CL is supported by Ghent University Geconcerteerde Onderzoeksactiviteiten (GOA) and Methusalem programs. The authors thank Ms. Astrid Luypaert and Dr. Viacheslav Mylka for assistance with the RNASeq set-up, Dr. Julie Deckers for assistance with the *in vivo* handlings, Dr. Rens M.J.M. de Vries for assistance with biophysical assays, Ms. Karima Bakkali for technical assistance with some of the reporter assays, Dr. Laurens Vyncke for advice on random mutagenesis analysis, Dr. Anastasia Kralli for sharing PGC1α plasmids, Dr. René Houtman for help with the interpretations of the peptide cofactor analyses, Dr. Sam Lievens and Dr. Irma Lemmens for their expert advice with MAPPITarray data analysis and Dr. Marnik Vuylsteke from GNOMIXX bvba, Statistics for Genomics, for assistance with statistical analyses. Dr. Lisa Koorneef is cordially thanked for the detailed proofreading of the content.

## Abbreviations

ERRα: estrogen-related receptor α
FAO: fatty acid β-oxidation
LBD: ligand-binding domain
MAPPIT: Mammalian Protein-Protein Interaction Trap
MARCoNI: Micro-Array for Real-time Coregulator and Nuclear receptor Interactions
NR: nuclear receptor
PDK4: pyruvate dehydrogenase linase 4
PGC1α: PPARγ coactivator 1 alpha
PPARα: Peroxisome-proliferator activated receptor α
PPRE: PPAR response element.

## Supplementary Figures

### Materials and methods unique to Supplementary data

#### Fluorescence polarization assay

A dilution series of XCT790 (100 µM – 48 pM) was made to a constant concentration of FL-ERRα (1 µM) and fluorescein-labeled PGC-1α cofactor peptide (residue 137-155; 0.1 µM) in a 384-well low volume black round bottom plate (Corning®). The buffer contained 25 mM Tris (pH=7.8), 200 mM NaCl, 5 mM DTT and 0.1% (w/v) bovine serum albumin. Plates were incubated for 30 minutes at 4°C before the fluorescence polarization was measured with a Tecan Infinite F500 plate reader using an excitation and emission wavelength of 485 nm and 535 nm respectively. The experiment was performed in duplicate and the data were analyzed using Prism (Graphpad).

### Supplementary data

**Supplementary Figure S1.**
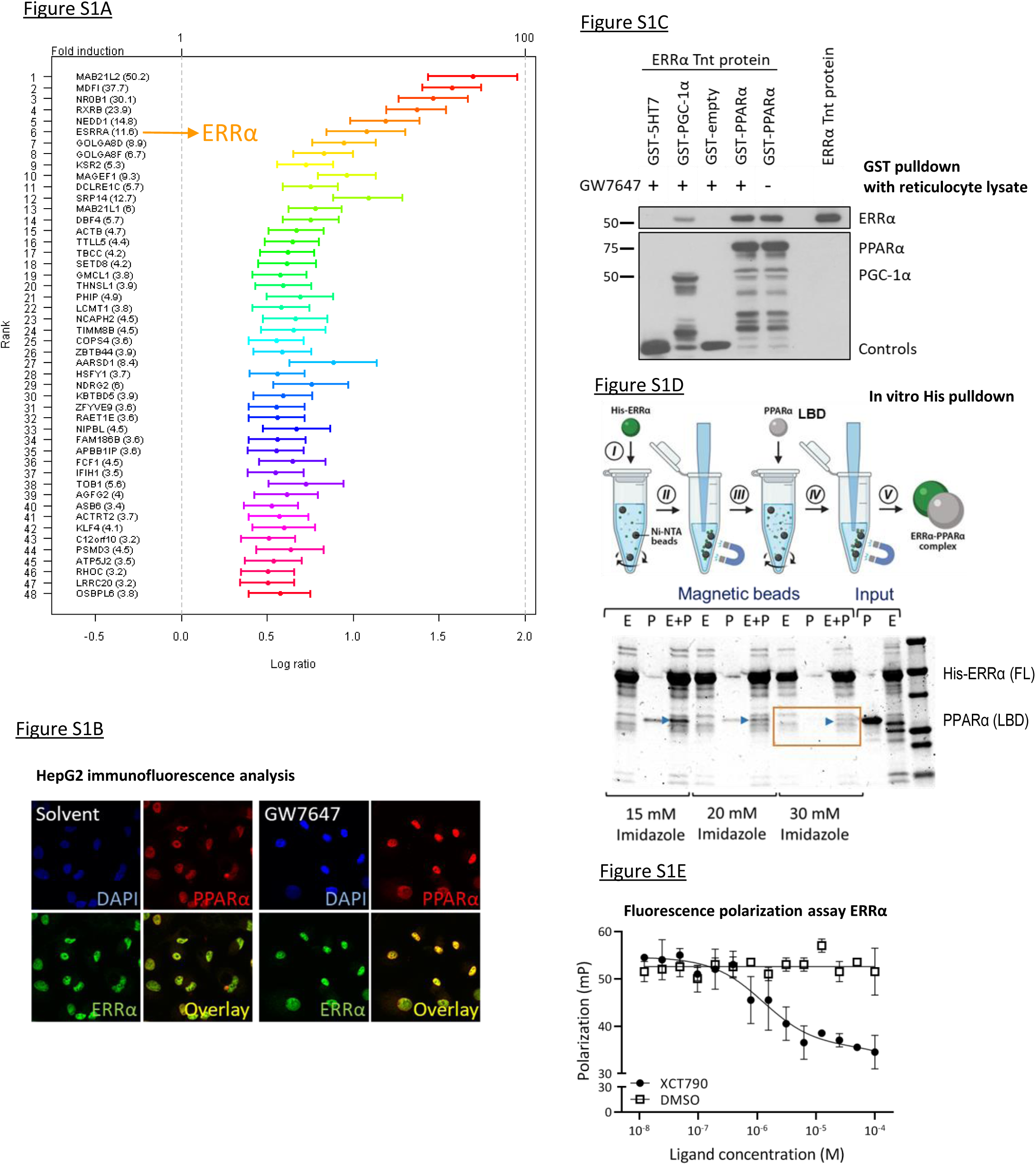
Characterization of the interaction between PPARα and ERRα in vitro and in cellulo. (A) Extended figure of Figure 1A. Array MAPPIT screen result of the activated (with GW7647) PPARα bait against a library covering up to 8500 preys, displayed as a ranking of the top PPARα interactors based on induction folds. (B) Serum-starved HepG2 cells were stimulated with GW7647 (0.5µM) for 1h. After fixation, cells were subjected to immunostaining with a mouse antibody against PPARα and a rabbit antibody against ERRα. The endogenous proteins were visualized with the secondary antibodies anti-mouse-Alexa 568 (red) and anti-rabbit-Alexa 488 (green). DAPI staining (blue) was used to visualize nuclei. (C) GST-pulldown between GST-PPARα and in vitro translated ERRα (ERRα TnT protein) using reticulocyte lysate. GST-5HT7 and GST-empty were used as negative controls, GST-PGC-1α as positive control. The PPARα ligand GW7647 was added in a high concentration (5µM) (n=3, representative figure). (D) His6-tag pulldown via FL-ERRα; Legend: E = FL-ERRα-His6, P = strep-PPARα-LBD (E) Fluorescence polarization assay. The ERRα-specific inverse agonist XCT790 was titrated to a constant concentration of bacterially expressed FL-ERRα with fluorescein-labeled PGC-1α peptide. Upon binding of the ligand, helix 12 of ERRα changes from the active conformation to an inactive conformation, releasing the PGC-1α cofactor.

**Supplementary Figure S2.**
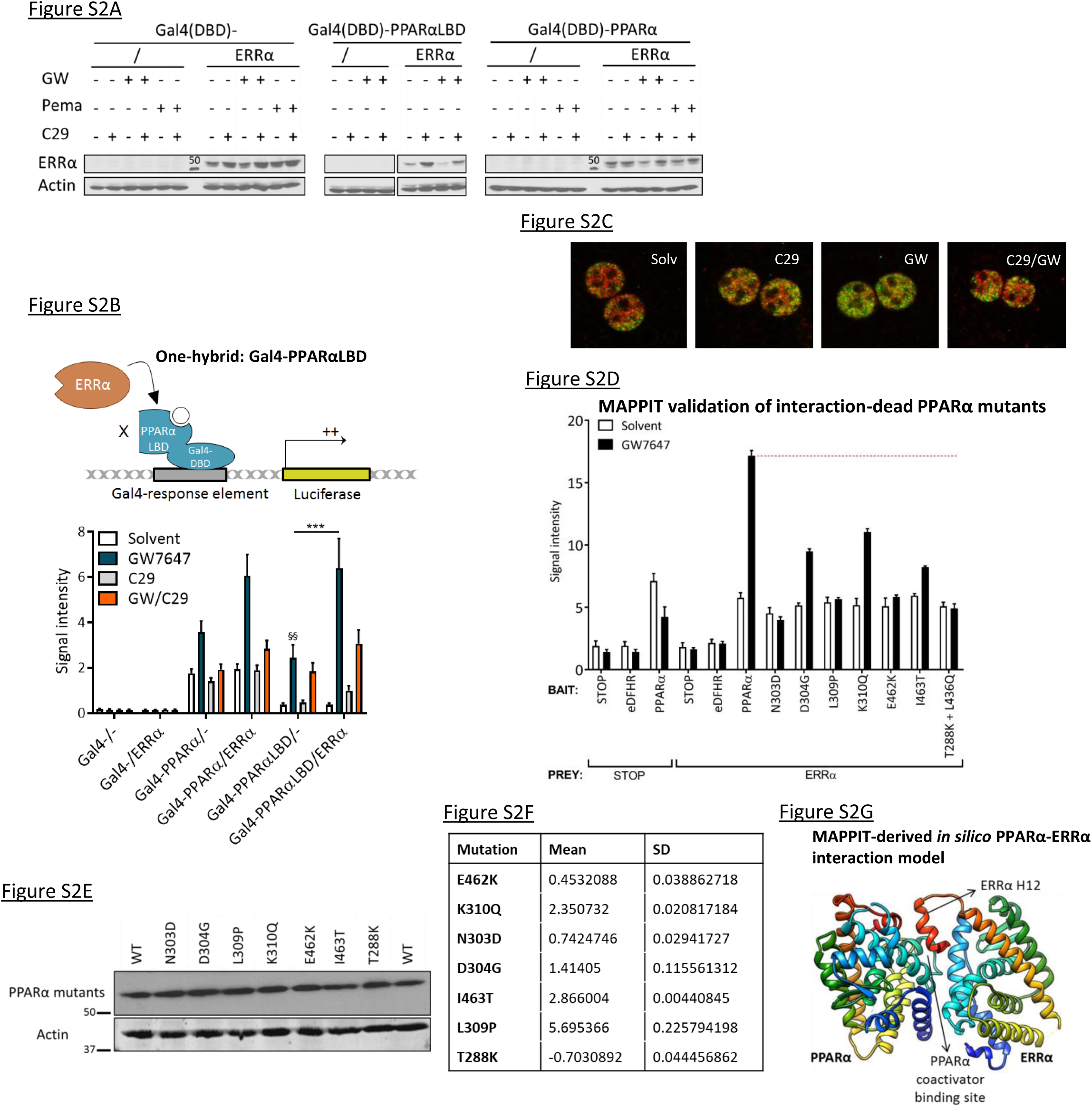
Impact of the pharmacological inhibition of ERRα and surface study of the PPARα-ERRα interaction. (A) Total cell lysates were prepared from a representative GAL4 experiment (see Fig. 2B, Fig. S2B) and subjected to Western Blot analysis. Actin served as a loading control, ERRα was detected via its Flag-tag. Abbreviations: GW, GW7647; pema, pemafibrate. (B) PPARαLBD (depicted in the model) is coupled to the DNA binding domain (DBD) of Gal4, which can bind its response element and activate the luciferase reporter. HepG2 cells, transfected with Gal4-responsive luciferase reporter, Gal4(DBD)-control or Gal4(DBD)-PPARαLBD, and with or without ERRα, were stimulated with GW7647 (GW, 0.5µM) and/or C29 (5µM) for 24h or were left untreated (Mean + SEM, n=3). In the Gal4-PPARαLBD conditions, the significance of differences in reporter activity was evaluated with unbalanced two-way ANOVA, followed by multiple comparison using the Fisher’s LSD test (*: p<0.05; **: p<0.01; ***: p<0.001, significance of single compound vs Solvent is marked with § sign). (C) GW7647 (0.5µM, 1h) -/+ C29 (5µM)-stimulated or Solvent control mouse primary hepatocytes were immunostained for PPARα (green) and ERRα (red); overlays are shown. (D) Validation of Fig. 2C. MAPPIT with PPARα wild-type (WT) or selected PPARαLBD mutants as bait and ERRα as prey. Empty prey (STOP), empty bait (STOP) and eDHFR-bait are used as negative controls. Cells were stimulated with leptin (100ng/ml) and leptin in combination with GW7647 (0.5µM) for 24h or were left untreated. Luciferase measurements were performed in triplicate (normalized by β-galactosidase expression) and normalized by untreated values. (E) HEK293T cells were transfected with a selection of PPARα-LBD mutants, corresponding to Fig. 2C. The PPARα WT plasmid was loaded as a positive control. Total cell lysates were prepared and subjected to Western Blot analysis. Actin served as a loading control. (F) The FoldX ΔΔg values, which correspond to the energy difference between WT and the indicated mutant, are listed for the selection of PPARα-LBD mutants with the X-ray structure of PPARα-LBD (PDB: 3VI8) as template. Each value is the average of five FoldX runs (Mean + SD). FoldX values higher than 1 indicate the possibility of disturbance by the mutation on protein folding/stability. (G) Theoretical and hypothetical molecular model for direct PPARα-ERRα interaction. The interaction involves the coactivator binding site of PPARα-LBD (PDB: 2P54) and helix 12 of ERRα-LBD (PDB: 3D24), which are marked on the ribbon structures.

**Supplementary Figure S3.**
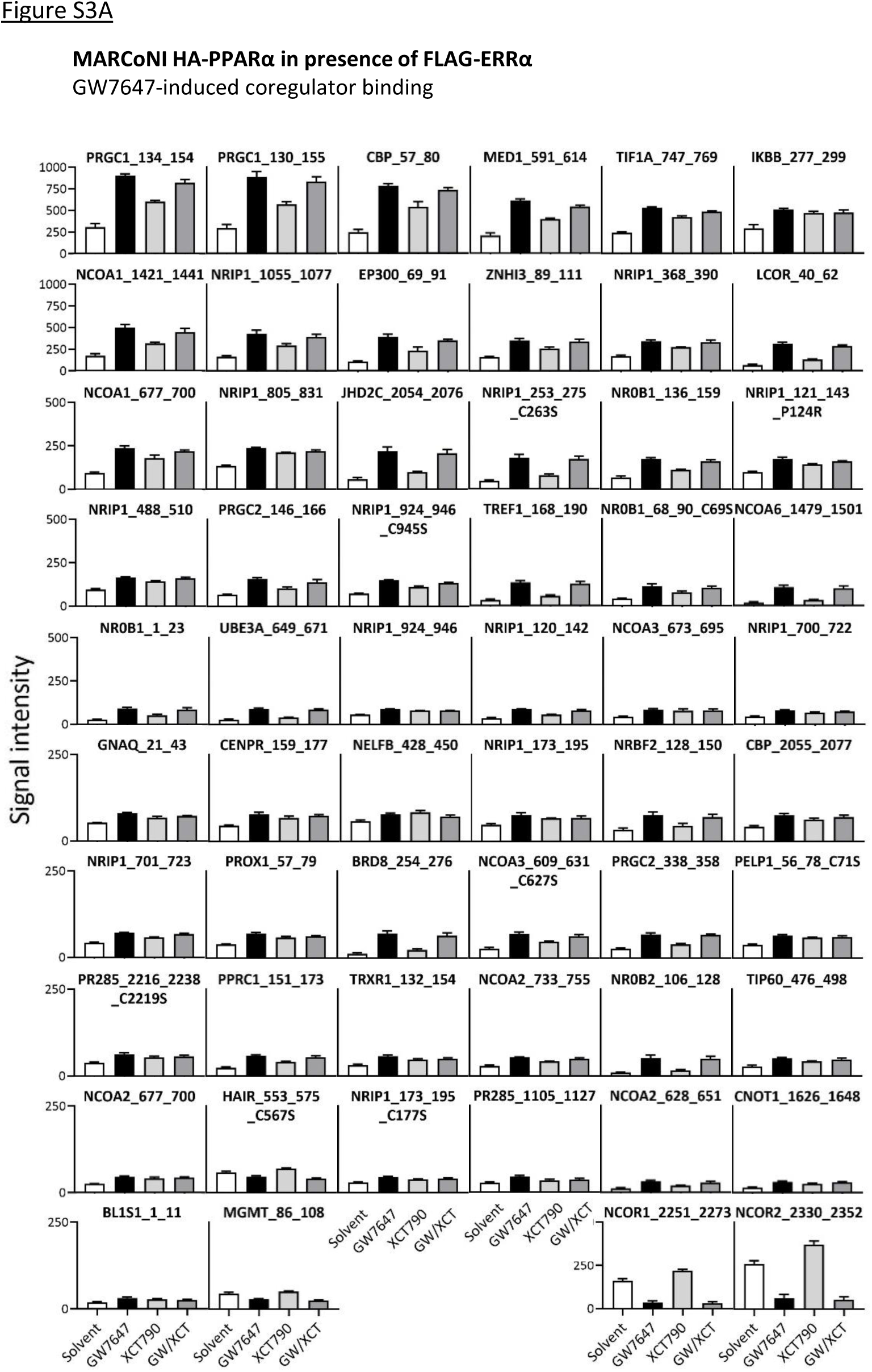
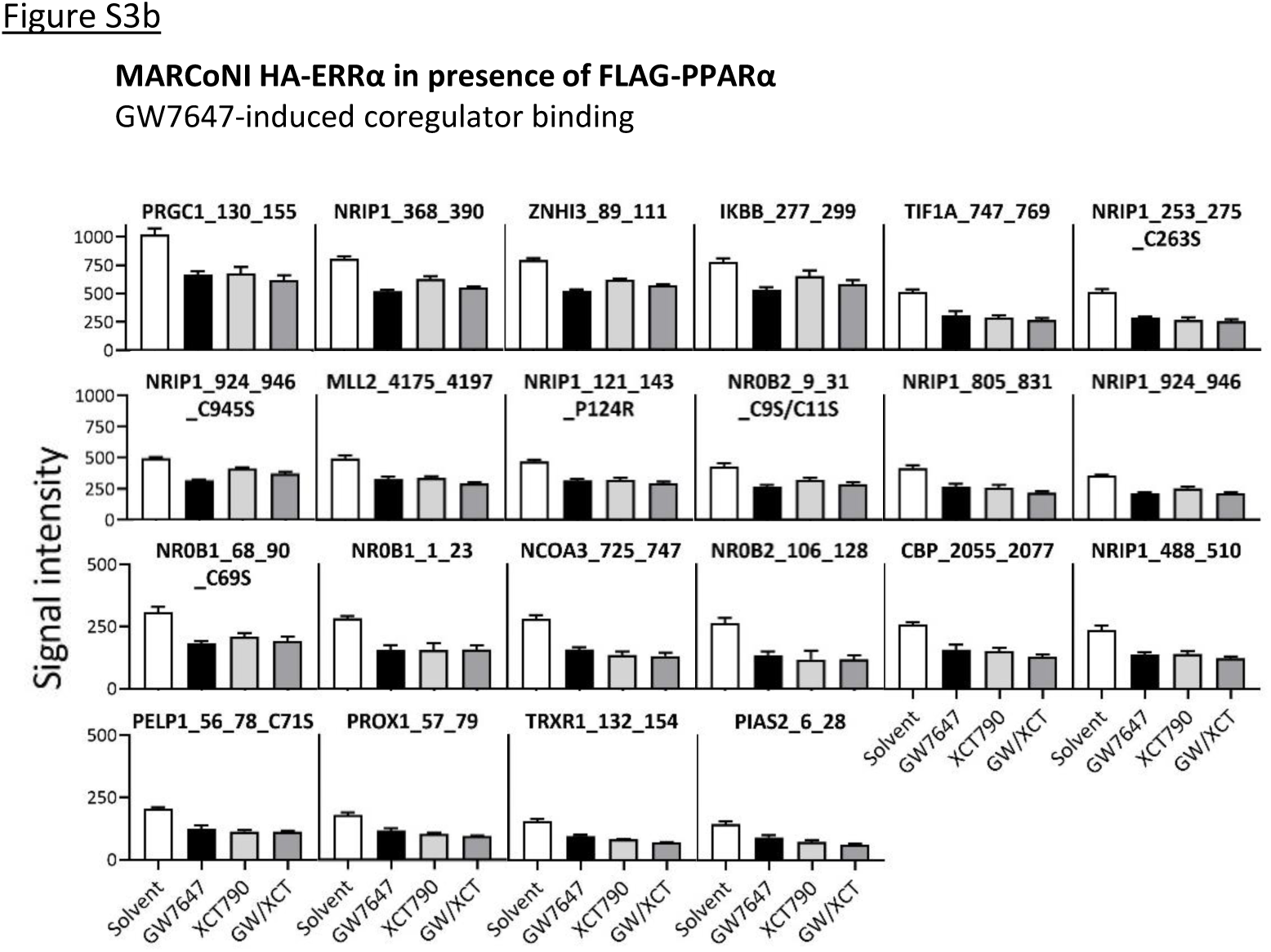
PPARα and ERRα cofactor profiling. (A) MARCoNI assay result with HA-PPARα, in the presence of FLAG-ERRα, overexpressed in HEK293T cells and stimulated with GW7647 (0.5µM) and/or XCT790 (1µM) for 2h. The significant binders upon GW7647 stimulation (Welch t-test, Benjamini-Hochberg p-value correction) are shown here, sorted upon signal intensity with GW7647 (with the two NCOR peptides at the end). Data was fitted according to a LOESS regression (Mean + SEM, n=3). (B) MARCoNI assay result with HA-ERRα, in the presence of FLAG-PPARα, overexpressed in HEK293T cells and stimulated with GW7647 (0.5µM) and/or XCT790 (1µM) for 2h. The significant binders upon GW7647 stimulation (Welch t-test, Benjamini-Hochberg p-value correction) are shown here, sorted upon signal intensity with Solvent. Data was fitted according to a LOESS regression (Mean + SEM, n=3).

**Supplementary Figure S4.**
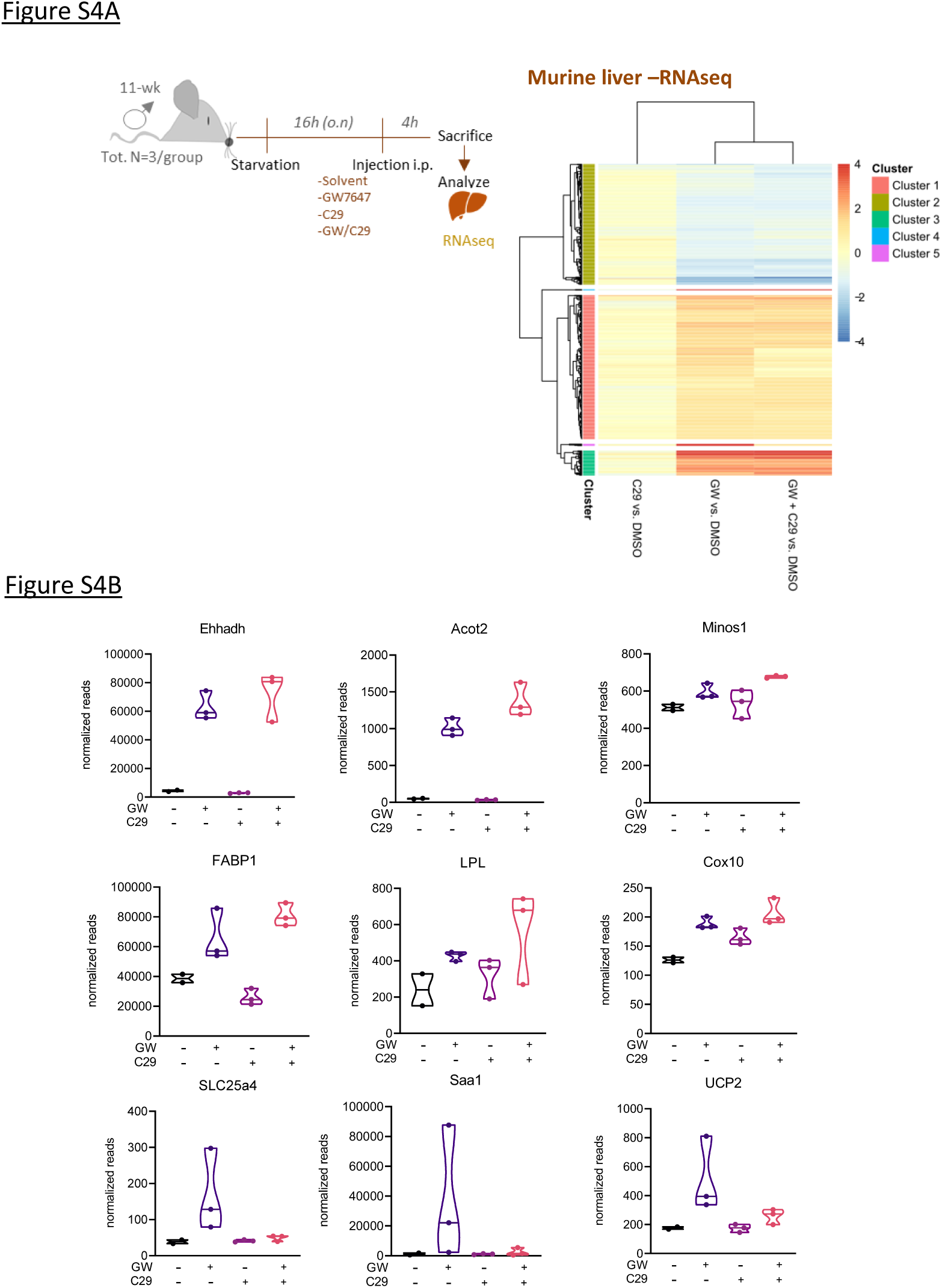
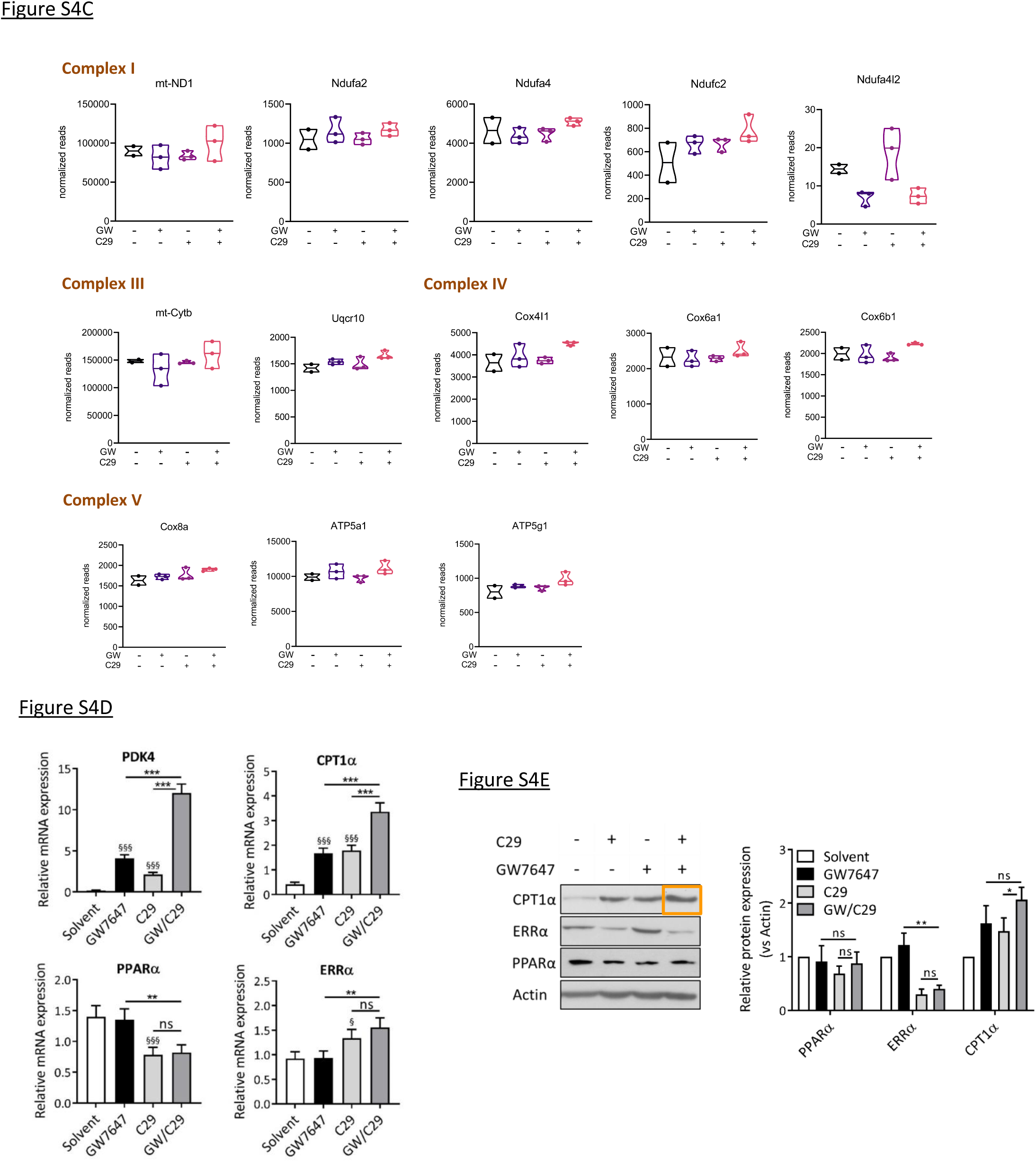
Genome-wide liver transcript analysis links crosstalk to fatty acid degradation and oxidative phosphorylation pathways. (A) RNAseq experiment with 3 mice/group followed by differential expression analysis. Log2 fold changes were compared, clustered, and presented as heatmaps. (B) Normalized reads of more GW-regulated transcripts showing similar or different regulation patterns. (C) Normalized reads showing the regulation of transcripts involved in the oxidative phosphorylation pathway. (D) Serum-starved HepG2 cells were stimulated with GW7647 (0.5µM) and/or C29 (5µM) for 24h. RNA expression values were normalized to the reference genes GAPDH and TBP using qBase+. Means + SE (n=6) are shown on the original scale. The significance of gene-specific ligand effects, estimated as differences (on the transformed scale) to the gene-specific reference level GW/C29, was assessed (*: p<0.05; **: p<0.01; ***: p<0.001, significance of single compound vs Solvent is marked with § signs). (E) Serum-starved HepG2 cells were stimulated with GW7647 (0.5µM) and/or C29 (5µM) for 24h. Total cell lysates were prepared and subjected to WB analysis (actin: loading ctrl). Protein expression values, obtained with Image J analysis of four independent replicates, were normalized to the loading control Actin and expressed as induction factor versus Solvent (Mean + SEM). The significance of differences in relative protein expression was evaluated with two-way ANOVA, followed by multiple comparison using the Fisher’s LSD test (*: p<0.05; **: p<0.01; ***: p<0.001).

**Supplementary Figure S5.**
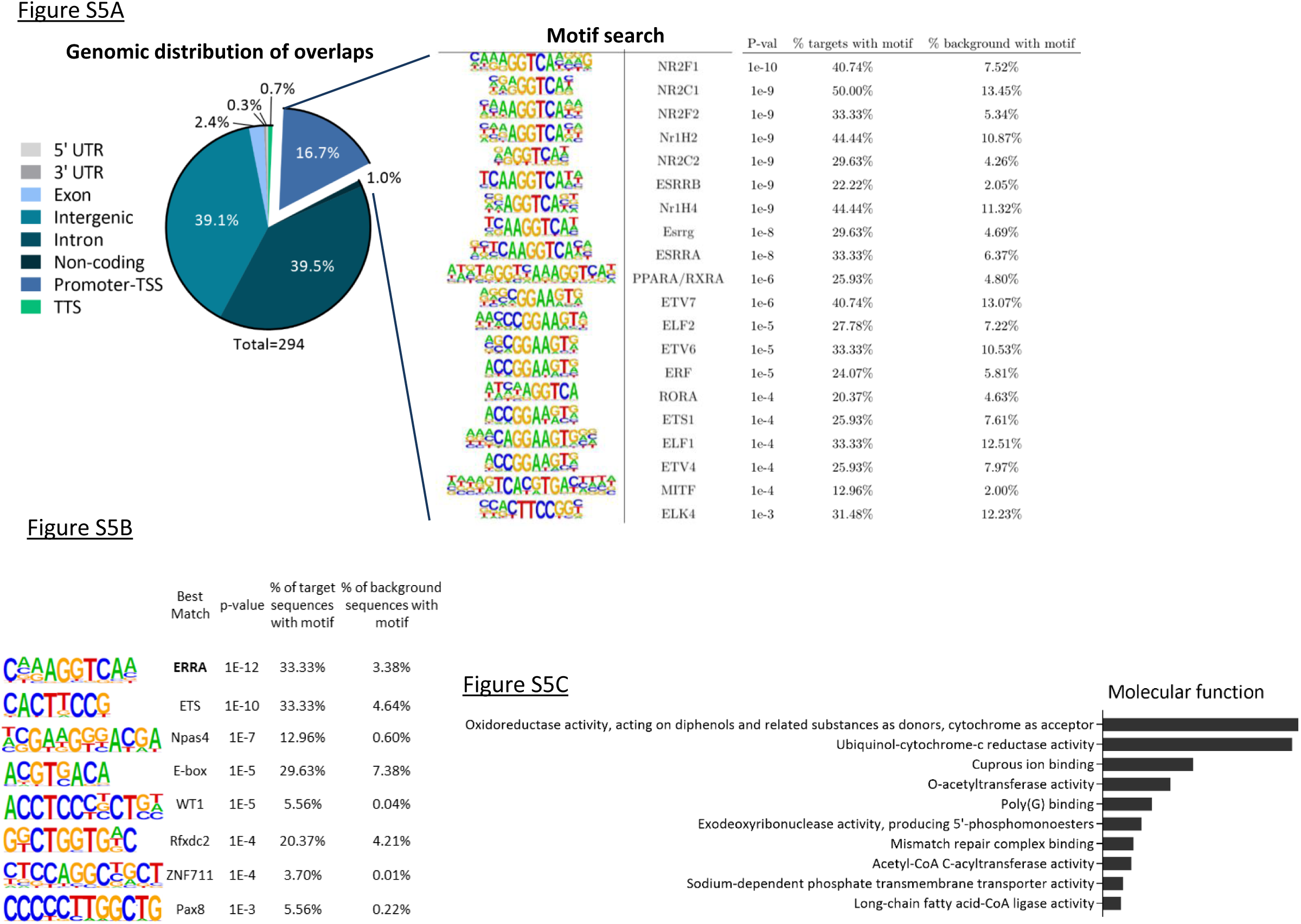
Genome-wide in silico cistrome analysis reveals overlapping PPARα and ERRα peaks and oxidative phosphorylation as a gene ontology enriched functionality. (A) Genomic distributions of the joint 294 overlapping (≥ 1bp) PPARα-ERRα peaks (see Figure 4G, 245+49 segments). Motif enrichment on the genes, located in the promoter-TSS region, annotated with the overlapping PPARα-ERRα peaks (49 peaks in total, Supplementary Table 1). The top 20 found motifs are shown here. (B) de novo motif enrichment on the genes, located in the promoter-TSS region, annotated with the overlapping PPARα-ERRα peaks (49 peaks in total, Supplementary Table 1). (C) gene ontology enrichment for ‘molecular function’. The categories are ranked according to p-value (lowest p-value on top).

**Supplementary Figure S6.**
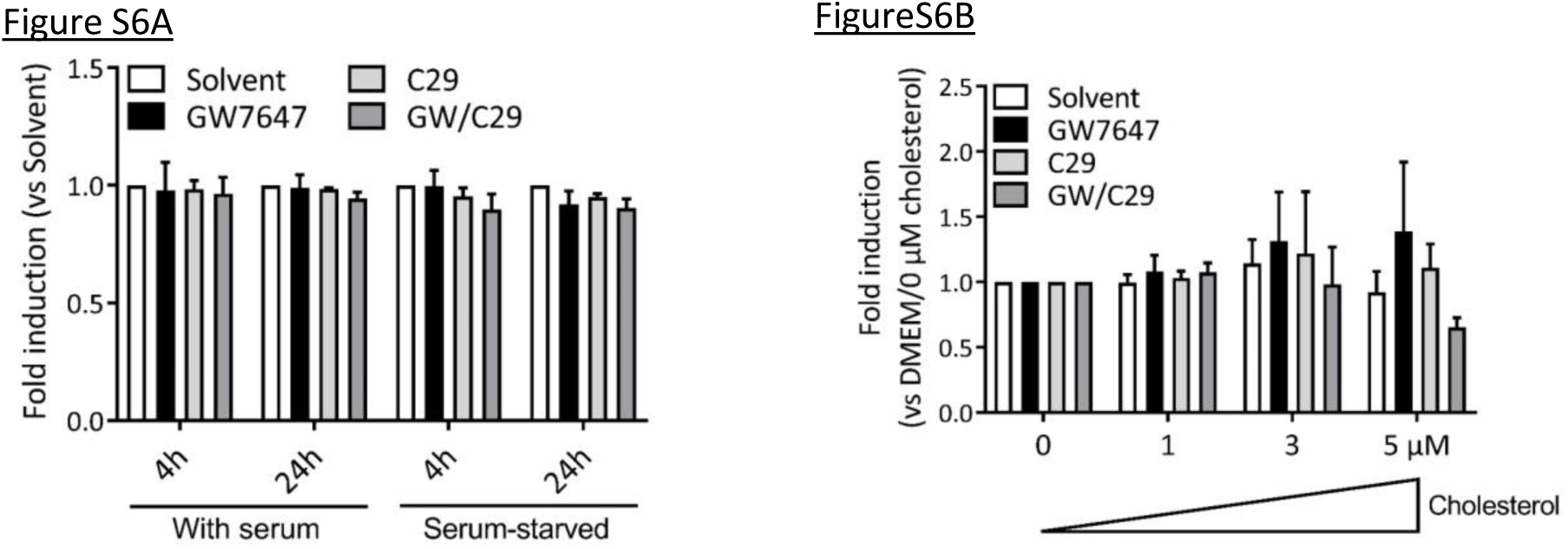
Serum-starvation, inductions and the cholesterol gradient do not affect cell viability. (A) and (B) For the analysis of cell viability, a Cell Titer-Glo luminescent cell viability kit was used in parallel to measure ATP production. Results are reported as fold induction versus Solvent in each condition, to evaluate the impact of the inductions and treatments on cell viability (Mean + SEM, n=3). The significance of differences in luminescence activity was evaluated with two-way ANOVA (none reached significance).

**Supplementary Figure S7.**
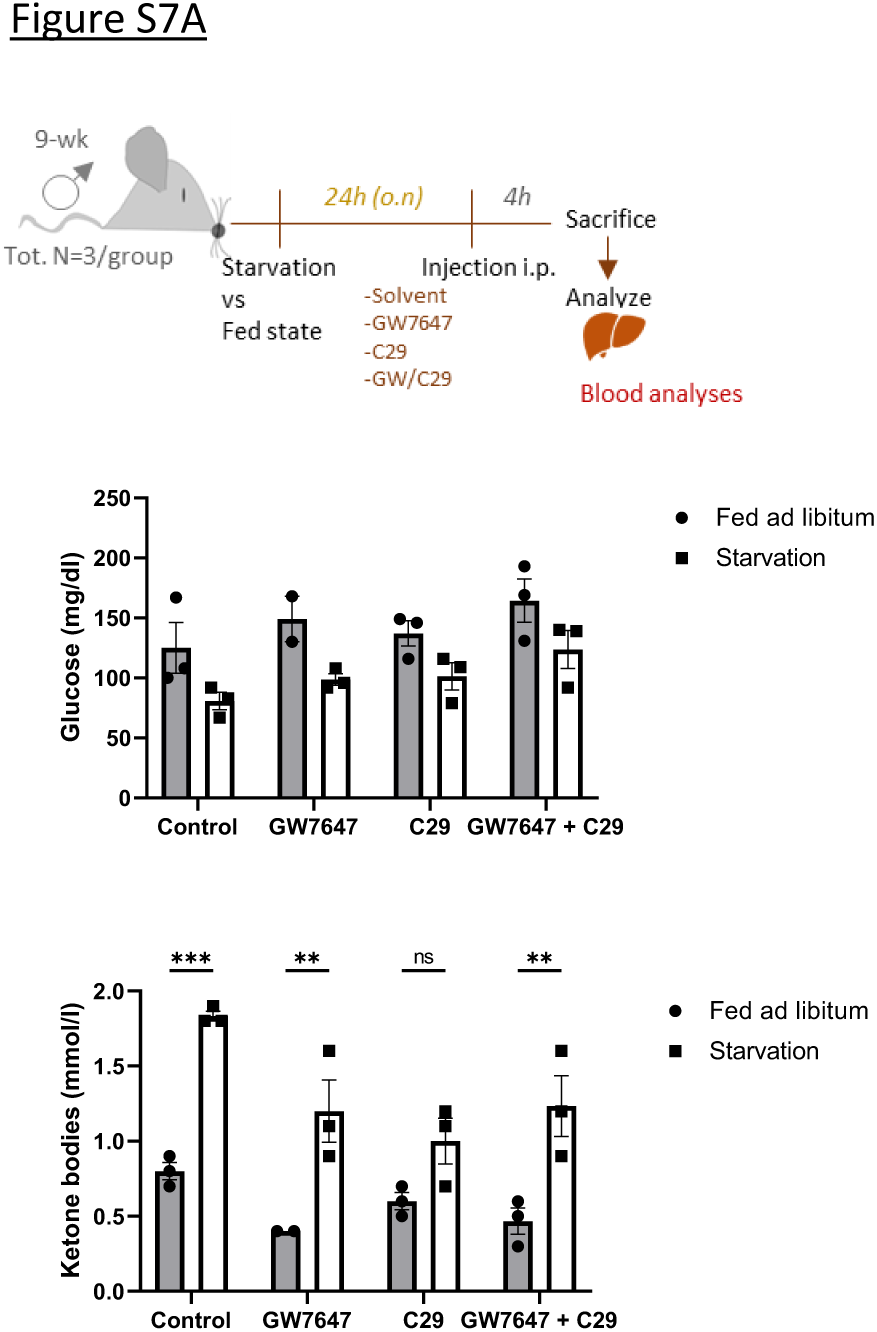
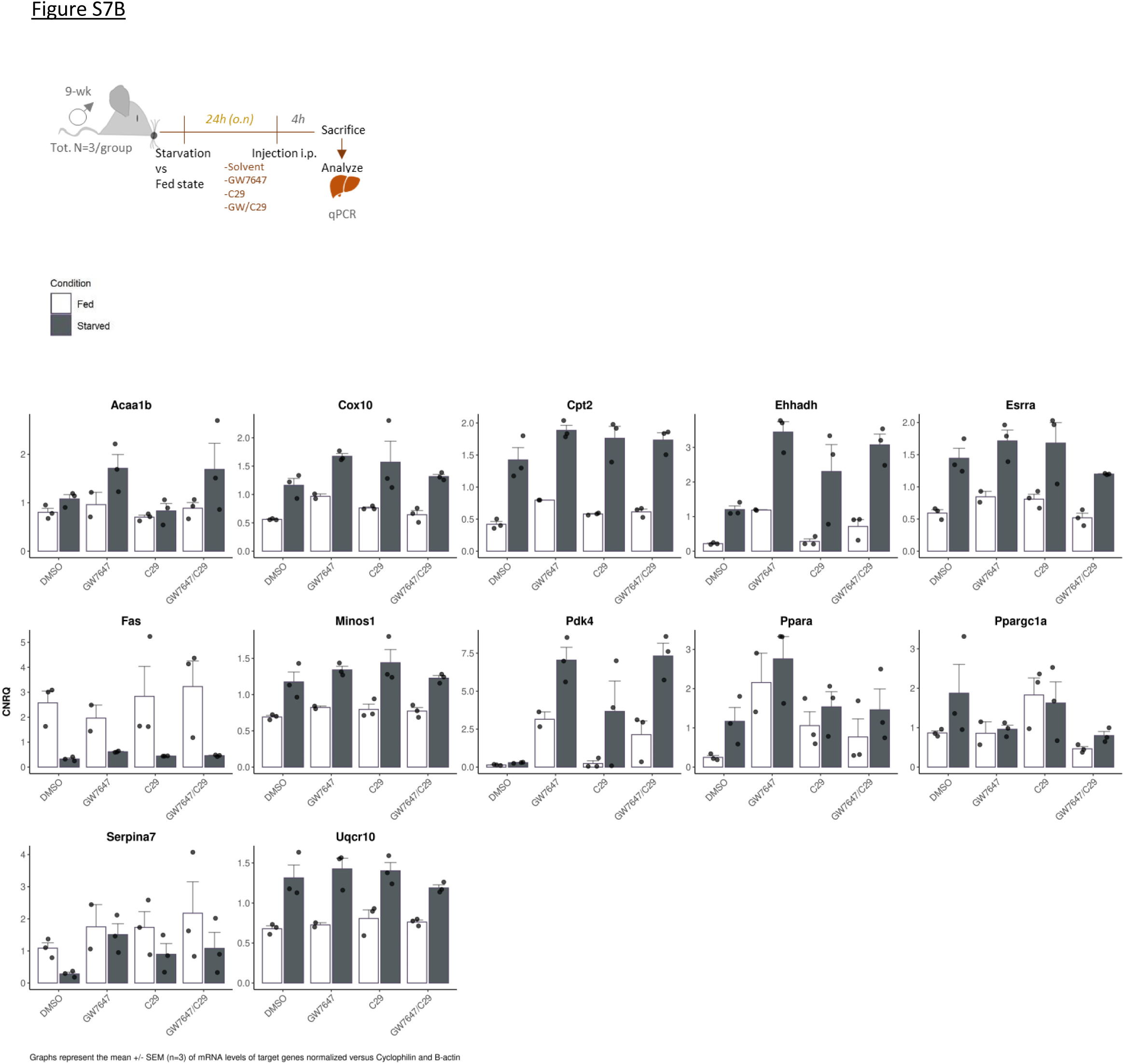
In vivo cross-talk study comparing livers subject to prolonged starvation versus fed states. (A) Male C57BI/6J mice (3 mice/group), after 24h starvation, were treated for 4h with GW7647 (4mg/kg) and/or C29 (10mg/kg) or were given solvent control, via i.p. injection. Blood glucose and ketone body levels were measured via tail blood. The significance of differences in treatment effect was evaluated using two-way ANOVA, followed by a multiple comparison analysis using the Sidak t-test (**: p<0.01; ***: p<0.001). (B) Male C57BI/6J mice, after 24h starvation or allowed feeding, were treated for 4h with GW7647 (4mg/kg) and/or C29 (10mg/kg) or were given solvent control, via i.p. injection. Livers were analyzed via qRT-PCR. RNA expression values were normalized to the reference genes Cyclophilin and b-actin using qBase+. Means +/- SEM (n=3).

**Supplementary Figure S8.**
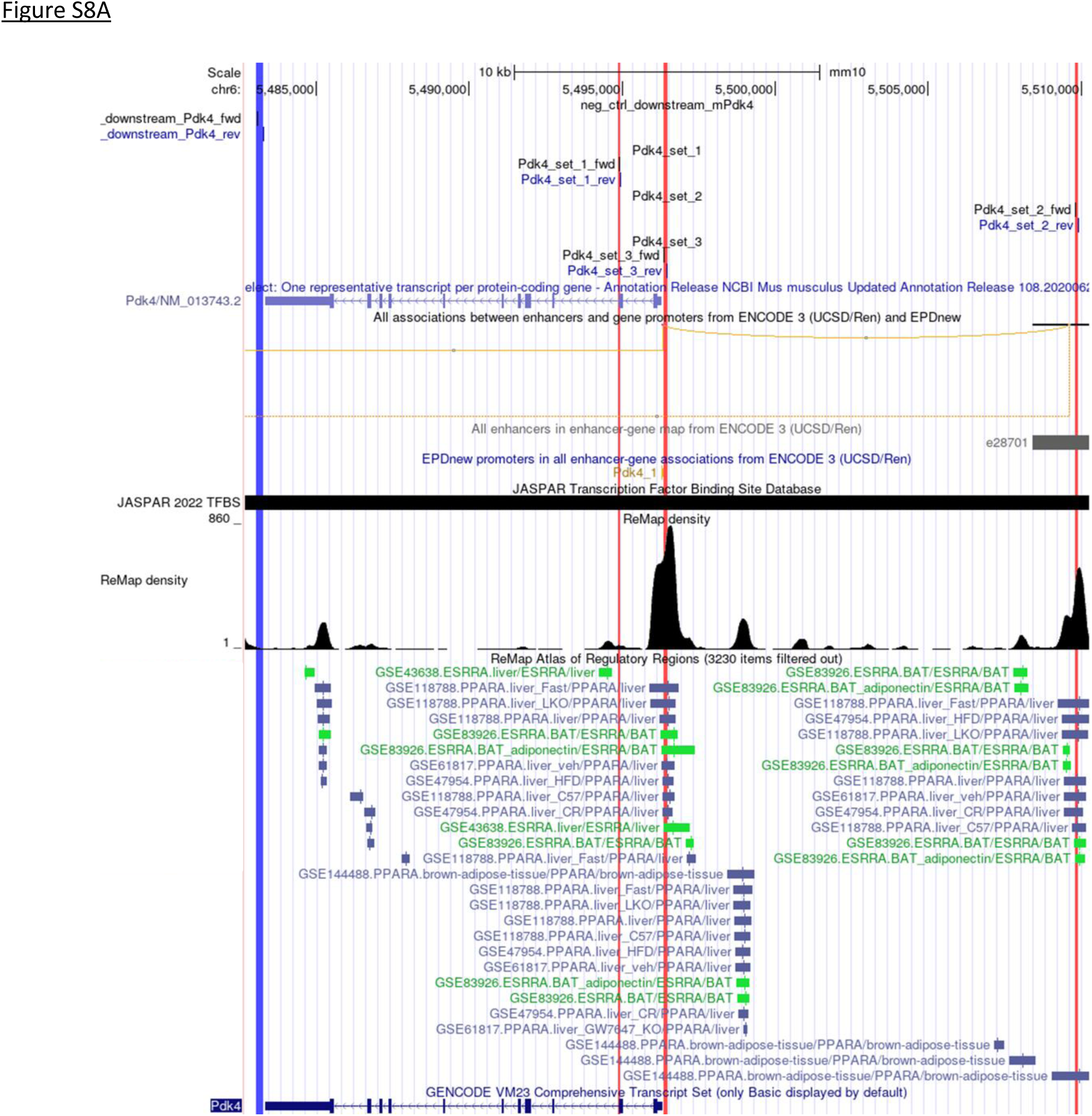
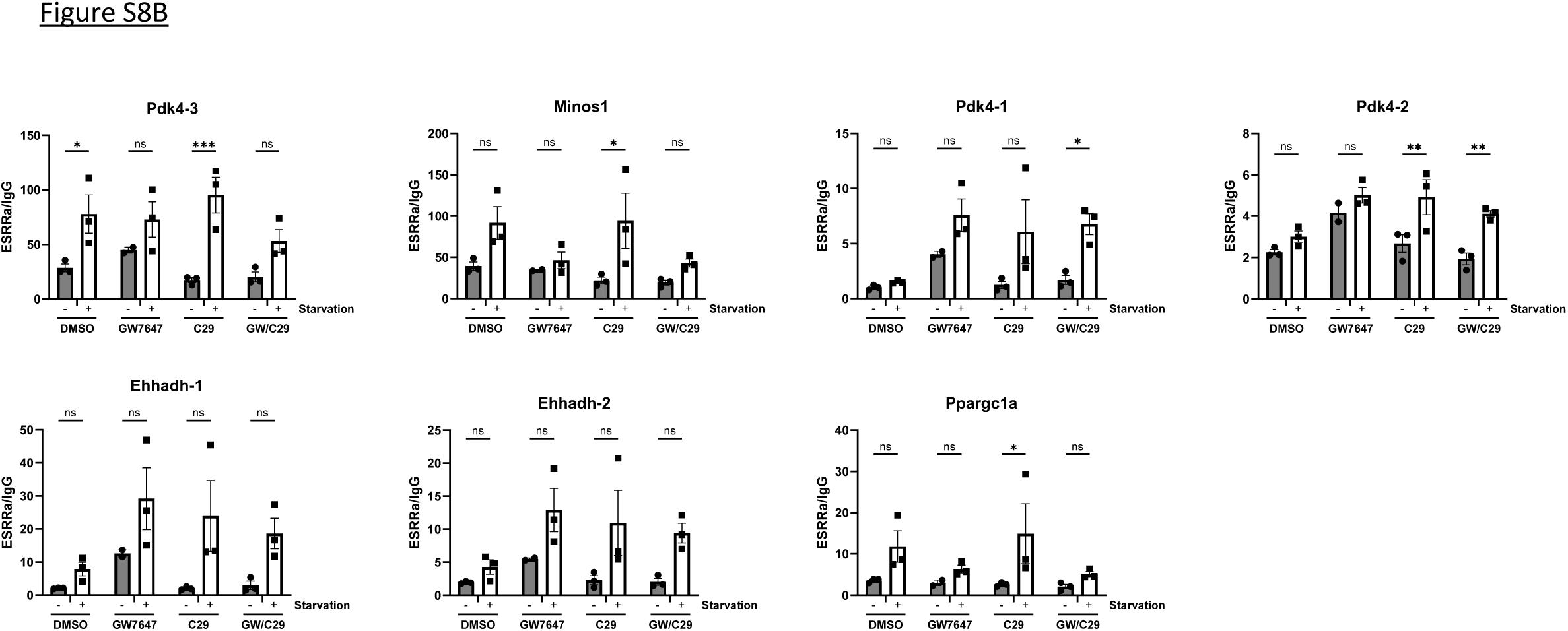
PPARα agonist treatment leads to ERRα recruitment at the chromatin of PPARα-dependent promoters and enhancers. (A) Localisations of the different ChIP primer sets in accord with control or in silico identified PPARα/ERRα binding regions for PDK4. Upstream region (blue line) or various promoter and enhancer regions (red lines) where PPARα (navy blue text) and/or ERRα (green text) binding is retrieved for PDK4 target gene regulations in various metabolic tissues. (B) C57BL/6J mice (n=3), after 24h starvation, were treated for 4h with GW7647 (4mg/kg) and/or C29 (10 mg/kg) or were given solvent control, via i.p. injection. Collected livers were used ChIP-qPCR which was performed with an anti-ERRα antibody versus IgG control. Results after immunoprecipitation were subtracted from the input and expressed as relative enrichment to the negative IgG control. The statistical significance was assessed using one-way ANOVA, followed by multiple comparison using the Tukey’s multiple comparison test (*: p<0.05; **: p<0.01; ***: p<0.001).

